# Single-cell analysis of pre-rRNA in *Escherichia coli* indicates distinct pathways of action for YbeX and YbeY proteins in ribosome biogenesis

**DOI:** 10.64898/2026.07.10.737703

**Authors:** Abbas Mansour, Ismail Sarigül, Tanel Tenson, Ülo Maiväli

## Abstract

The *Escherichia coli* protein YbeX/CorC is encoded in the same operon as the ribosome biogenesis factor YbeY, and its deletion leads to accumulation of 17S pre-rRNA and degradation intermediates of 16S rRNA under magnesium limitation. To further investigate the *ybeX* deletion phenotype, we used rRNA fluorescence *in situ* hybridization coupled with flow cytometry (rRNA-FISH-flow) to quantify 16S rRNA, 23S rRNA, and 17S pre-rRNA levels at single-cell resolution in *ΔybeX* and *ΔybeY* strains. *ΔybeX* cells grown under limiting Mg^2+^ develop striking cell-to-cell heterogeneity in 17S pre-rRNA content during the transition to stationary phase, with up to 25-fold differences between individual cells. Upon regrowth from the stationary phase, *ΔybeX* cultures display a bimodal distribution of 17S pre-rRNA, revealing two distinct subpopulations — one retaining high levels of unprocessed pre-rRNA and the other with low levels — whose relative proportions shift over time, until visible growth resumes. The stoichiometry between mature 16S and 23S rRNAs remains tight in both strains, indicating that the heterogeneity is specific to pre-rRNA processing, rather than a general disruption of ribosome homeostasis. The *ΔybeY* mutant accumulates 17S pre-rRNA more uniformly across cells and primarily during exponential growth in rich medium, consistent with its direct role in 16S rRNA maturation. These single-cell data suggest that YbeX and YbeY affect ribosomal RNA metabolism through distinct mechanisms and that the extended lag phase of *ΔybeX* is caused by a heterogeneous clearing of pre-ribosomal intermediates in individual cells.

## INTRODUCTION

Magnesium ion, the most abundant divalent cation in living cells, is essential for multiple cellular processes, and diverse transport systems maintain its homeostasis across all domains of life (Flatman 1991; Groisman et al. 2013). These transporter systems enable adaptation to fluctuations in Mg^2+^ availability, promote bacterial virulence in response to Mg^2+^ limitations, while supporting cellular growth and fundamental processes by maintaining adequate intracellular Mg^2+^ (Groisman and Chan 2021; Franken et al. 2022).

As Mg^2+^ is essential for ribosome biogenesis, maintaining its homeostasis is needed for ribosome assembly and efficient protein synthesis (Akanuma 2021). Within the ribosome, Mg^2+^ promotes the association of the small and large subunits, stabilizes the secondary structure of rRNA, and facilitates the binding of ribosomal proteins (Klein et al. 2004). When extracellular Mg^2+^ levels decrease, bacterial cells tend to lower intracellular Mg^2+^, thus reducing ribosome content and inhibiting growth (Pontes et al. 2016). Upon Mg^2+^ starvation, *Escherichia coli* intact ribosomes dissociate into inactive smaller subunits with lower sedimentation values (S) that subsequently undergo further degradation (McCarthy 1962; Maiväli et al. 2013).

CorC domain-containing proteins are widely distributed in all domains of life and are crucial for maintaining ionic balance and Mg^2+^ homeostasis in *Salmonella* and *E. coli* (Sarıgül et al. 2024; Iwadate and Slauch 2025). YbeX/CorC in *E. coli* exhibits 97% sequence identity to the *Salmonella* homolog and is encoded within the same operon as the ribosome-associated proteins YbeY and YbeZ. The three proteins are members of the σ^32^ heat shock regulon (Nonaka et al. 2006). YbeY is involved in ribosome biogenesis, RNA processing, quality control, and turnover, whereas YbeZ interacts with YbeY and multiple ribosomal proteins (Andrews and Patrick 2022; Butland et al. 2005; Liao et al. 2021; Summer et al. 2020). While the co-encoding of YbeX suggests a potential functional link to ribosome biogenesis, the specific role of this protein remains unknown.

We previously showed that the *E. coli ybeX* knockout strain (*ΔybeX*) exhibits a prolonged lag phase upon regrowth, specifically when cells are pre-grown under low-Mg^2+^ conditions (Sarıgül et al. 2024). The mutant exhibits heterogeneous phenotypes regarding colony size and increased sensitivity to elevated temperature and ribosome-targeting antibiotics that induce cold shock responses. *ΔybeX* cells accumulate pre-16S rRNA (17S rRNA) and 16S rRNA degradation intermediates, and these phenotypes are rescued by Mg^2+^ supplementation, supporting the notion that CorC regulates ribosomal metabolism through magnesium homeostasis (Sarıgül et al. 2024).

Given that the *ΔybeX* cells exhibited heterogeneous phenotypes at the population level (as indicated by heterogeneous colony sizes), we investigated the effect of *ybeX* knockout on ribosome abundance under Mg^2+^ limitation at single-cell resolution using rRNA fluorescence *in situ* hybridization (rRNA-FISH). We also characterized the *ΔybeY* phenotype to compare the effects of the two deletions on ribosome metabolism. Our data show that *ybeX* knockout cells exhibit pronounced alterations in rRNA processing in the stationary phase, with a substantial accumulation of 17S rRNA under low Mg^2+^ conditions. Although both *ybeX* and *ybeY* knockouts affect ribosome metabolism, their effects on rRNA levels across growth stages differ markedly.

## RESULTS

### Experimental setup

We previously showed that *ΔybeX* cells grown into the late exponential phase under limiting Mg^2+^ display a prolonged lag phase upon outgrowth in limiting Mg^2+^ (Sarıgül et al. 2024). This phenotype coincides with a gradual accumulation of 17S pre-rRNA during growth under low Mg^2+^. We used 50 µM Mg^2+^ as the minimum concentration that allows bacterial growth (Sarıgül et al. 2024). Once growth resumes, the growth rates of *ΔybeX* and isogenic wild-type (WT) cultures are indistinguishable. These observations led us to hypothesize that accumulation of inactive, 17S-containing pre-ribosomes underlies the extended lag phase of *ΔybeX*. Here, we test this conjecture by quantifying rRNA species at the single-cell level.

Isogenic *E. coli* BW25113 (WT) and *ΔybeX* strains were inoculated into MOPS medium containing 10 mM MgCl₂, grown overnight, and then diluted into fresh Mg^2+^ -limited MOPS (50 µM MgCl₂). Samples were collected at the time points indicated in Figure 1a. A part of each culture was incubated overnight (15 h), then diluted again into fresh medium containing 50 µM MgCl₂ and grown for a further 12 h with regular sampling. The two rounds of growth were thus designed to (i) accumulate 17S pre-rRNA and prime the outgrowth phenotype in the first round, and (ii) quantify the clearance of 17S pre-rRNA during subsequent outgrowth in the second round.

**Figure 1.**
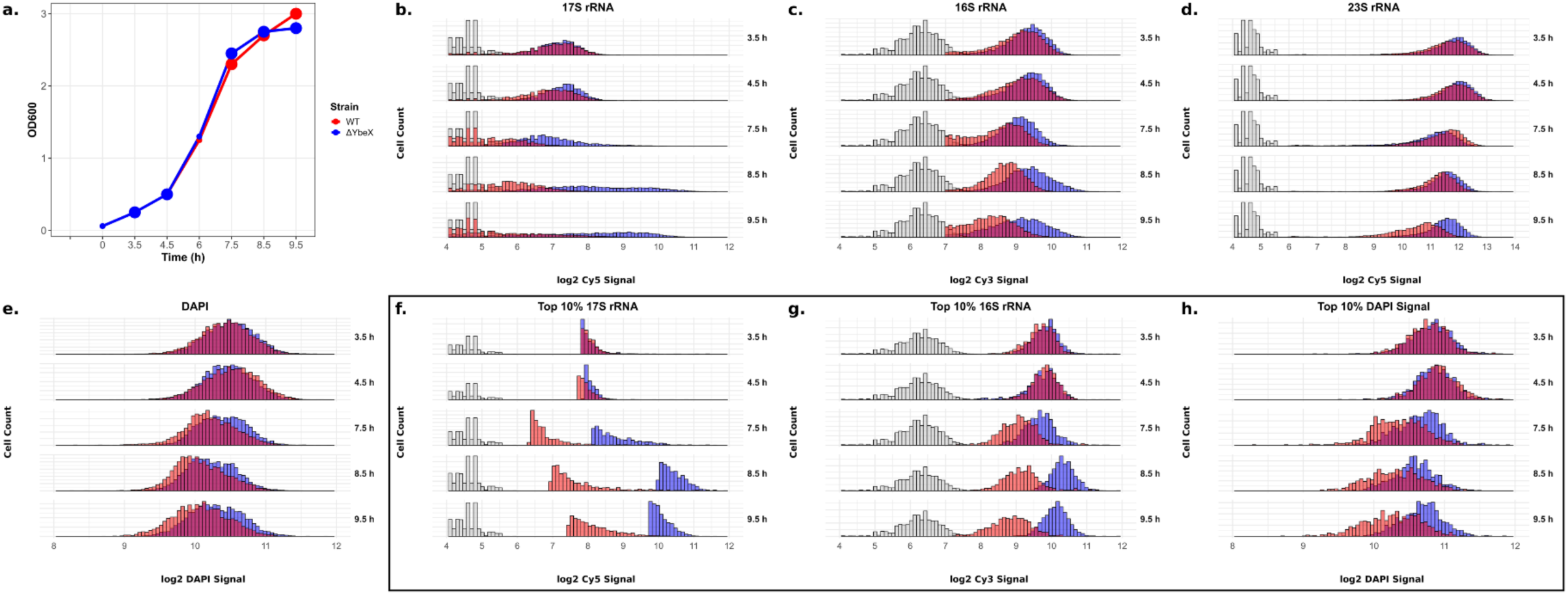
Per-cell rRNA distributions in WT and ΔybeX in the first growth cycle under Mg^2+^ limitation. (a) Growth curves at OD_600_ for isogenic E. coli BW25113 (WT, red) and ΔybeX (blue) cultures grown in Mg²⁺-limited MOPS medium following dilution from a high-Mg²⁺ overnight pre-culture. Enlarged points mark the sampling points shown in panels b–h. (b–d) Per-cell distributions of (b) 17S pre-rRNA, (c) 16S rRNA, and (d) 23S rRNA signals for WT (red) and ΔybeX (blue). X-axes show log2 fluorescence intensity. Gray histograms show the controls labeled with non-hybridizing fluorophores. (f, g) Signal from cells in the top 10% of the 17S pre-rRNA distribution within each sample, showing (f) 17S pre-rRNA and (g) 16S rRNA for the same subset of cells. (e) DAPI signal (DNA content) for the full cell population. (h) DAPI signal for the top 10% 17S subpopulation (i.e. the same cells shown in f and g). 23S and 17S signals are measured in Cy5; 16S signal is measured in Cy3. Distributions are of a representative experiment of at least three biological replicates; aggregate statistics across the replicates are reported in Supplementary Figure 2.

Each culture sample was divided into two subsamples, fixed, permeabilized, and hybridized with Cy3- and Cy5-labeled DNA oligonucleotide probes. One subsample received Cy5-anti-17S and Cy3-anti-16S probes; the other received Cy5-anti-23S and Cy3-anti-16S probes. It must be noted that the 16S rRNA-specific probe also binds 17S pre-rRNA, so that the nominal 16S signal is a composite of 16S and 17S contributions. Because 16S rRNA is far more abundant than 17S pre-rRNA in most samples, the composite signal is dominated by mature 16S rRNA; this interpretation is consistent with the strong correlation observed between the 23S and 16S rRNA signals (see below). We therefore treat the 16S rRNA probe output as reflecting mature 16S rRNA. DNA was stained with DAPI, and DAPI-negative particles were excluded from downstream analysis. DAPI signal distributions for the full cell population and the top 10% 17S pre-rRNA subpopulation are shown in Figures 1e and 1h, respectively.

### Ribosomal RNA phenotypes at the single-cell level

In the first round of growth, WT and *ΔybeX* cultures diluted from high-Mg^2+^ MOPS medium grew with similar kinetics during the exponential phase (Figure 1a, Supplementary Figure 1). The distributions of mature 23S and 16S rRNA were also similar between strains during early-to-mid exponential growth (Figure 1c, d). During mid-exponential growth, the mature-rRNA distributions of both strains shifted toward approximately 1.5-fold lower signals, with no substantial change in distribution width or shape. At the final time point, corresponding to the late-exponential/transition, the *ΔybeX* mature-rRNA distributions remained essentially unchanged while the WT distributions shifted further toward lower values. At this time point, the median *ΔybeX* cell carried roughly 2-fold higher mature rRNA signal than the median WT cell. Across strains and time points, the cell-to-cell spread of mature rRNA signal spanned approximately 10- to 15-fold (*ca*. 4 log₂ units; Figure 1c, d).

In the early-exponential phase, the 17S pre-rRNA distribution resembled that of mature rRNA, albeit at a median signal roughly 20-fold lower than 23S rRNA (assuming comparable probe-binding efficiencies; Figure 1b). From mid-exponential phase onward, WT 17S pre-rRNA levels declined in step with the slowing of growth and new rRNA synthesis. In *ΔybeX*, by contrast, median 17S levels rose during mid- to late-exponential growth. The increase was seen in most, but not all, cells and was accompanied by a marked broadening of the 17S distribution — cell-to-cell variation grew from approximately 5-fold to approximately 25-fold (≈4.6 log₂ units). At the upper tail of the distribution, *ΔybeX* cells reached 17S pre-rRNA levels approaching those of the lowest 23S rRNA signals, though not necessarily within the same cells.

These single-experiment observations were consistent with the aggregate summary across replicates, in which each sample was characterized by its median (central tendency) and interquartile range (cell-to-cell variation) before linear regression against time (Supplementary Figure 2). The aggregate analysis confirmed a rise in median 17S pre-rRNA levels in *ΔybeX* while WT medians stagnated. Both mature rRNA species were present at modestly higher levels (up to ∼1.5-fold) in *ΔybeX* across the growth curve.

### Reanalysis of the subpopulation of cells with the highest 17S pre-rRNA content

We next focused on the cell subpopulation with the highest 17S pre-rRNA levels, reasoning that very high 17S pre-rRNA levels in later growth phases are most likely to compromise mature rRNA levels and disturb rRNA stoichiometry in the *ΔybeX* strain (Sarıgül et al. 2024). For each sample, we reanalyzed the top 10% of cells by 17S pre-rRNA signal. In *ΔybeX*, this signal peaked at the penultimate 8.5-hour time point, reaching approximately one third of the 23S rRNA level; in WT, the same subpopulation carried 17S pre-rRNA roughly 20- to 30-fold below 23S rRNA (Figure 1f). At the final time point, the *ΔybeX* top 10% 17S pre-rRNA signal decreased approximately 1.5-fold within one hour of growth.

During growth into the late-exponential phase, WT cells showed a roughly two-fold drop in 16S and 23S rRNAs, whereas *ΔybeX* cells did not, leaving *ΔybeX* with *ca.* two-fold higher mature-rRNA concentrations (Figure 1c, d). This divergence was evident in both the full-data and top 10% 16S rRNA analyses (Figure 1g). During these final hours of growth — during which *ΔybeX* cells fail to down-regulate mature rRNA — the *ΔybeX* culture arrested (Figure 1a). We found no evidence for disrupted 16S–23S or 17S–16S rRNA stoichiometry in *ΔybeX* relative to WT (Supplementary Figure 3).

### The stoichiometry of the rRNA species

To assess stoichiometry between mature rRNA species and between 17S pre-rRNA and mature 16S rRNA, we computed Pearson correlation coefficients from the single-cell data for each sample and modeled their temporal evolution by linear regression (Figure 2). In a parallel analysis, we recalculated these coefficients for the top 10% of cells by 17S pre-rRNA signal from the Cy5-anti-17S/Cy3-anti-16S probe pair. We acknowledge that correlations computed over restricted signal ranges are mathematically biased toward lower values; nonetheless, our aim was to determine whether cells with extreme 17S pre-RNA levels differ from average cells in rRNA stoichiometry.

**Figure 2.**
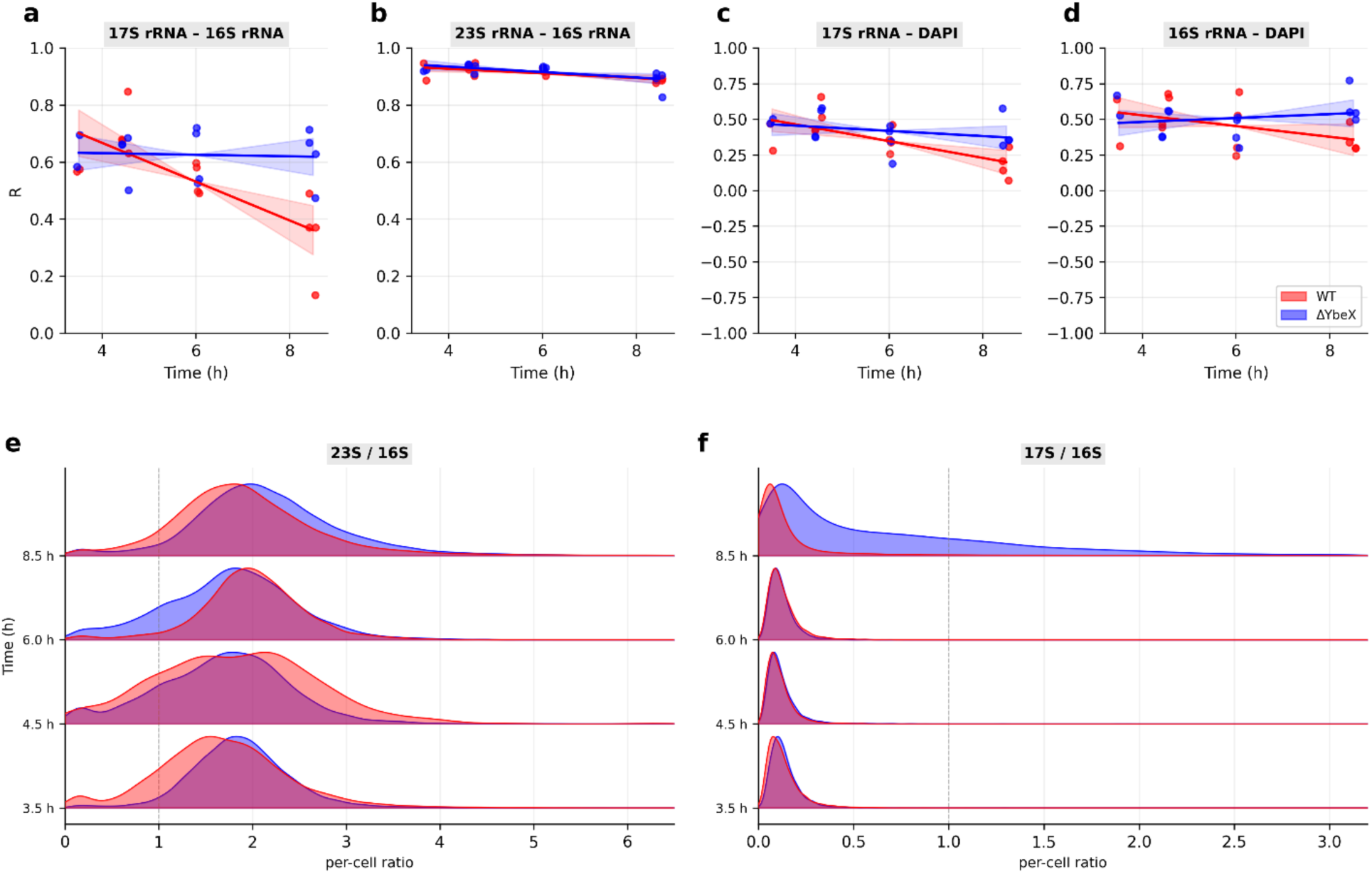
a) The stoichiometry of the rRNA species as assessed by Pearson correlation coefficients (R) of 17S pre-rRNA vs. 16S rRNAs, b) 16S vs. 23S rRNAs, c) 17S pre-rRNA-DAPI, d) 16S rRNA-DAPI, obtained from the full data. Temporal dynamics were modeled by linear regression. Shaded areas show 95% CI. Data points from independent experiments are represented by points (red for WT and blue for ybeX). e) Per-cell rRNA-ratio distributions (linear scale, dashed line = ratio of 1) for 23S/16S and f) 17S/16S ratios for WT (red) and ΔybeX (blue), time points shown as ridgelines. Individual experiments and the pooled composite are in Supplementary Figures 4 and 5.

In both WT and *ΔybeX*, we observed excellent correlation, and therefore stoichiometry, between mature 16S and 23S rRNAs. This correlation was highest in the exponential phase and declined only slightly by the end of the growth cycle. Cell-to-cell correlations between 17S and 16S rRNAs were substantially weaker than mature-rRNA correlations, averaging ∼0.65 in *ΔybeX* and falling from ∼0.7 to ∼0.35 in WT as cultures progressed from early to late exponential phase (Figure 2). This strain difference is plausibly a consequence of the strong reduction in WT 17S pre-rRNA levels at later time points: in the top-10% 17S subpopulation, we observed the opposite trend — an increase in 17S–16S rRNA correlation (Supplementary Figure 3).

Whereas a correlation between two rRNA species reflects subunit stoichiometry, a correlation between an rRNA species and DNA content (DAPI) can reflect the coupling between ribosome content, translational capacity, and cell division. We observed a moderate correlation between mature 16S rRNA and DAPI signal (R² ≈ 0.15). The corresponding correlation for 17S pre-rRNA was substantially weaker (Figure 2).

We also plotted individual cell-level ratios of 23S/16S and 17S/16S rRNAs (Figure 2e,f, Supplementary Figures 4, 5). Because the ratios are of cy5/cy3 signals attached to DNA probes with different (and unknown) binding efficiencies to different rRNAs, we have no way of knowing which exact ratio would correspond to 1:1 stoichiometry. However, the ratios relative to WT/mutant strains are informative about both potential differences in mean ratios and the ranges of cell-to-cell variation under different conditions.

At the single-cell level, the 23S/16S rRNA ratio distributions of WT and *ΔybeX* overlapped closely at every time point of the first growth cycle (median linear ratio ≈ 1.4–1.7 in both strains), and the cell-to-cell spread of this ratio stayed constant throughout (10th–90th-percentile fold-range ≈ 1.6–2.5), consistently with tight 16S–23S rRNA stoichiometry. The 17S/16S rRNA ratio behaved differently. While WT and *ΔybeX* were indistinguishable during early-to-mid exponential growth (3.5–6 h; median 17S/16S ≈ 0.20–0.23, fold-range ≈ 2.5–3 in both strains), at the final, late-exponential time point (8.5 h), the *ΔybeX* 17S/16S rRNA ratio distribution broadened: its 10th–90th-percentile fold-range expanded to ≈ 8.5-fold (against ≈ 5.6-fold in WT), from ≈ 2.5–3-fold at earlier time points (Figure 2f). Thus, the single-cell ratio distribution confirms that mature-subunit (23S/16S rRNA) stoichiometry remains tightly balanced in *ΔybeX*, while the divergence in 17S/16S rRNA stoichiometry at the transition to stationary phase is driven by a widening of the cell-to-cell distribution rather than by a uniform shift.

### 17S rRNA accumulated from stationary phase is heterogeneous among cells

We next analyzed the second round of the experiment, in which stationary-phase cultures from the first round were diluted into fresh Mg²⁺-limited medium. We reproduced the previously reported extended lag phase of *ΔybeX* followed by WT-like outgrowth rates (Figure 3a; Supplementary Figure 1; (Sarıgül et al. 2024)). Immediately after dilution, the signals of both mature rRNA species were approximately two-fold lower in WT than in *ΔybeX*. This difference disappeared after one hour of growth, a time point at which neither culture had yet resumed visible growth. For the following four hours, the WT and *ΔybeX* mature-rRNA distributions were very similar, even as WT entered early exponential growth and *ΔybeX* remained in lag. As soon as *ΔybeX* resumed visible growth, its mature rRNA levels rose, re-establishing a ∼two-fold difference to WT by the final time point. In parallel, the subpopulation of *ΔybeX* cells with very high 17S pre-rRNA signal gradually disappeared, so that by the time *ΔybeX* had completed its first few divisions, the 17S profiles of the two strains were essentially indistinguishable (Figure 3a, b).

**Figure 3.**
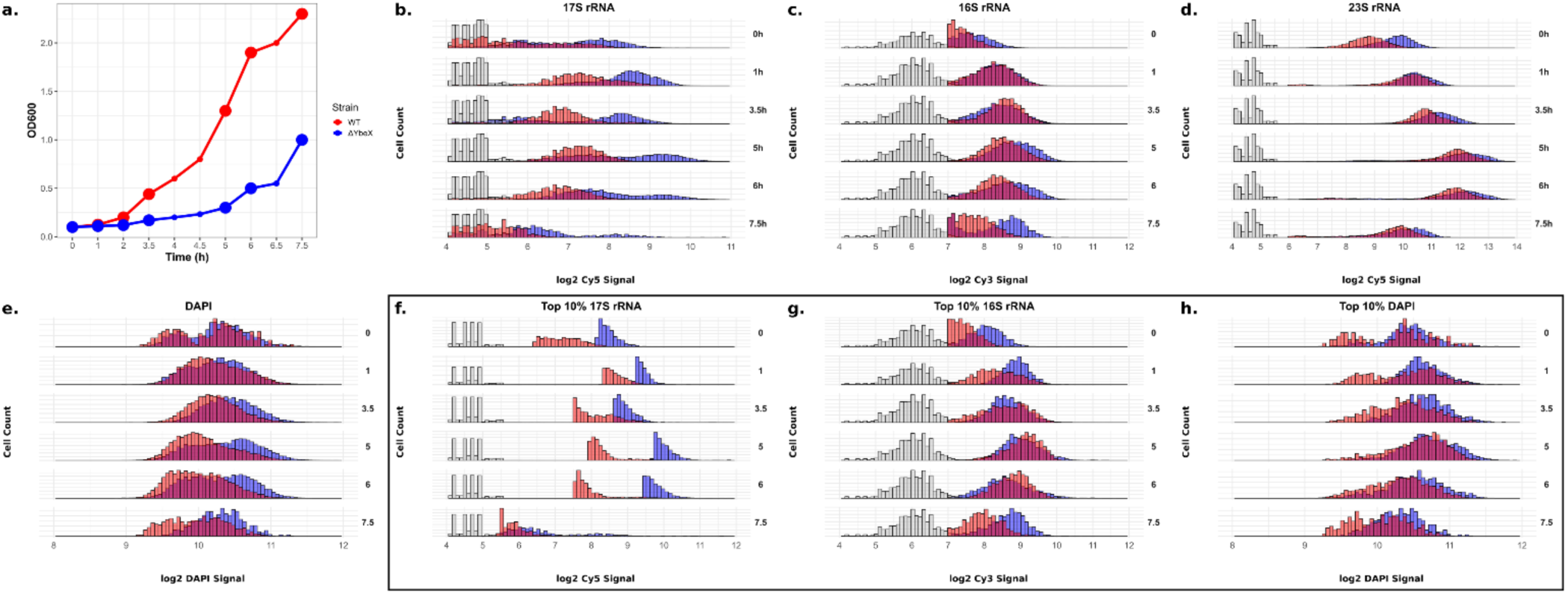
The outgrowth experiment involves growing cells in a medium with limiting MgCl_2_ until they reach the stationary phase (the 1^st^ growth cycle), then diluting them into fresh medium and allowing them to enter their second exponential phase in a medium with limiting MgCl_2_ (the 2^nd^ growth cycle). a. Full data of the 2^nd^ growth cycle experiment, b. Top 10% subpopulation. Analysis of a representative experiment is presented as in Figure 1.

Unlike the first round, in the second round the *ΔybeX* 17S pre-rRNA distribution during lag was distinctly bimodal, indicating two coexisting subpopulations with qualitatively different pre-ribosome states (Supplementary Figure 6). The relative proportions of the two subpopulations shifted over time toward the low-17S mode, and visible growth resumed when the two modes were approximately equally populated (∼5 h). A two-peaked distribution was also visible in the DAPI signal of both WT and *ΔybeX*, with a modestly higher mean in *ΔybeX* (Figure 3). As in the first growth round, the correlation between 17S pre-rRNA and DAPI was weak, especially in the top-10% 17S pre-rRNA subpopulation (Figure 4; Supplementary Figure 10).

**Figure 4.**
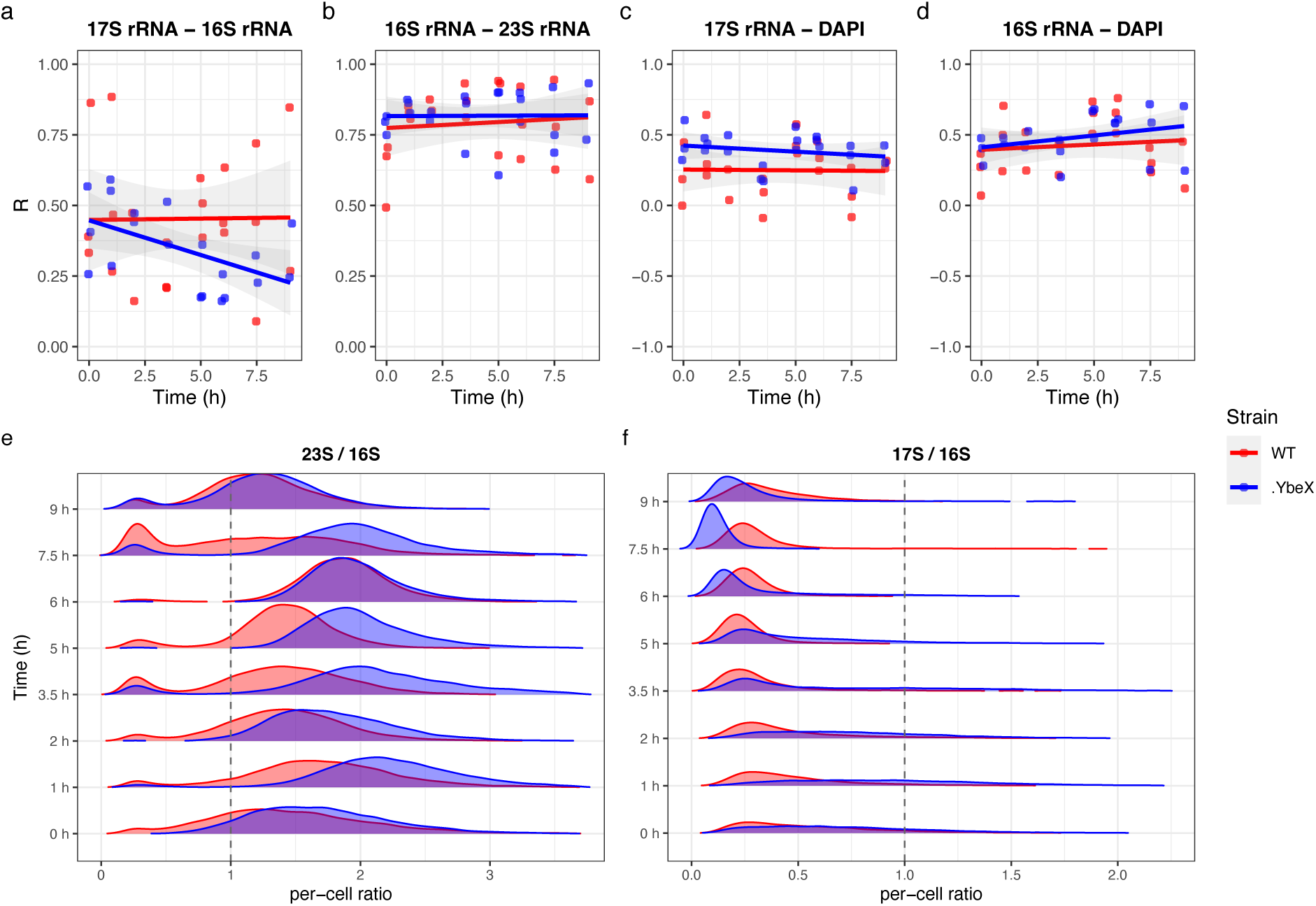
The stoichiometry of a) 17S vs. 16S rRNAs and b) 16S vs. 23S rRNAs, c) 17S vs DAPI d) 16S vs DAPI from the second growth round full data, assessed by linear correlation. For each time point and strain in each experiment, Pearson correlation coefficients (R) were calculated, and temporal dynamics were modeled by pooled linear regression. Data points from independent experiments are represented by points (red for WT and blue for ΔybeX) and regression lines with shaded 95% CI are colored accordingly. e) Per-cell rRNA-ratio distributions for 23S/16S and f) 17S/16S ratios for WT (red) and ΔybeX (blue) across the second-cycle time points. Individual experiments and the pooled composite are in Supplementary Figures 7 and 8.

The bulk-level correlations between both 16S vs 23S and 17S vs rRNAs were somewhat lower in the second-round experiment, although the higher experiment-to-experiment variation in the second round calls for cautious interpretation of these results (Figure 4). However, the possibility of lower stoichiometry between mature rRNAs in the second growth round is supported by single-cell analysis of 16S-23S rRNA ratios, which show a much wider distribution in the second growth round than in the first (Fig 4 vs Fig 2). In both 16S-23S and 17S-16S distributions, we see initial narrowing upon resumption of growth, followed by re-widening as the cultures enter mid- and late exponential growth.

During the extended lag (0–3.5 h), the *ΔybeX* 17S/16S rRNA ratio was elevated roughly 2- to 3-fold over WT and showed a very wide cell-to-cell spread (10th–90th-percentile fold-range ≈ 4–6, rising to ≈ 10–15-fold at 6–7.5 h), consistent with the bimodal 17S pre rRNA distribution (Supplementary Figure 6). As *ΔybeX* resumed visible growth and cleared its accumulated pre-rRNA, the 17S/16S ratio fell to the WT value by 7.5–9 h. (Figure 4e,f, replicates and pooled data are shown in Supplementary Figures 7 and 8).

The aggregate analysis of the second growth cycle showed that, as growth resumed, median mature rRNA rose and median 17S pre-rRNA fell. As in round one, all rRNA species were present at higher levels in *ΔybeX* than in WT (Supplementary Figure 9). Unlike in round one, DAPI signal was also slightly higher in *ΔybeX*.

### The YbeY knockout accumulates rRNA during early growth

To place the *ΔybeX* results in a broader context, we asked how the single-cell rRNA landscape of *ybeY*, a second ribosome-associated deletion mutant from the same operon, compares with that of *ΔybeX*. YbeX and YbeY are both associated with ribosome biogenesis and translation, but their phenotypes differ markedly. Unlike *ΔybeX*, *ΔybeY* has a strong constitutive growth defect that does not require Mg²⁺ limitation to manifest (Davies et al. 2010; Jacob et al. 2013) and is therefore readily studied in rich medium without the sensitizing conditions used for *ΔybeX*. Accordingly, we grew WT and *ΔybeY* in LB (rather than Mg²⁺-limited MOPS) and omitted the outgrowth experiment from the stationary phase, as there is no Mg²⁺-dependent priming step to exploit. This simpler single-cycle design mirrors the conditions under which the biochemical role of YbeY in 16S 3ʹ-end maturation was originally characterized.

WT and *ΔybeY* cultures grown in LB reached their highest 16S and 23S rRNA signals during mid-exponential growth (Figure 5a; OD₆₀₀ = 0.9–1.75). *ΔybeY* carried approximately 1.3-to 2-fold higher 16S and 23S signals than WT across this phase. As cultures entered the stationary phase, the mature rRNA signal declined roughly 3- to 4-fold in both strains. Both strains showed cell-to-cell variation of approximately 8-fold between the lowest- and highest-signal cells at every time point (Figure 5c, d).

**Figure 5.**
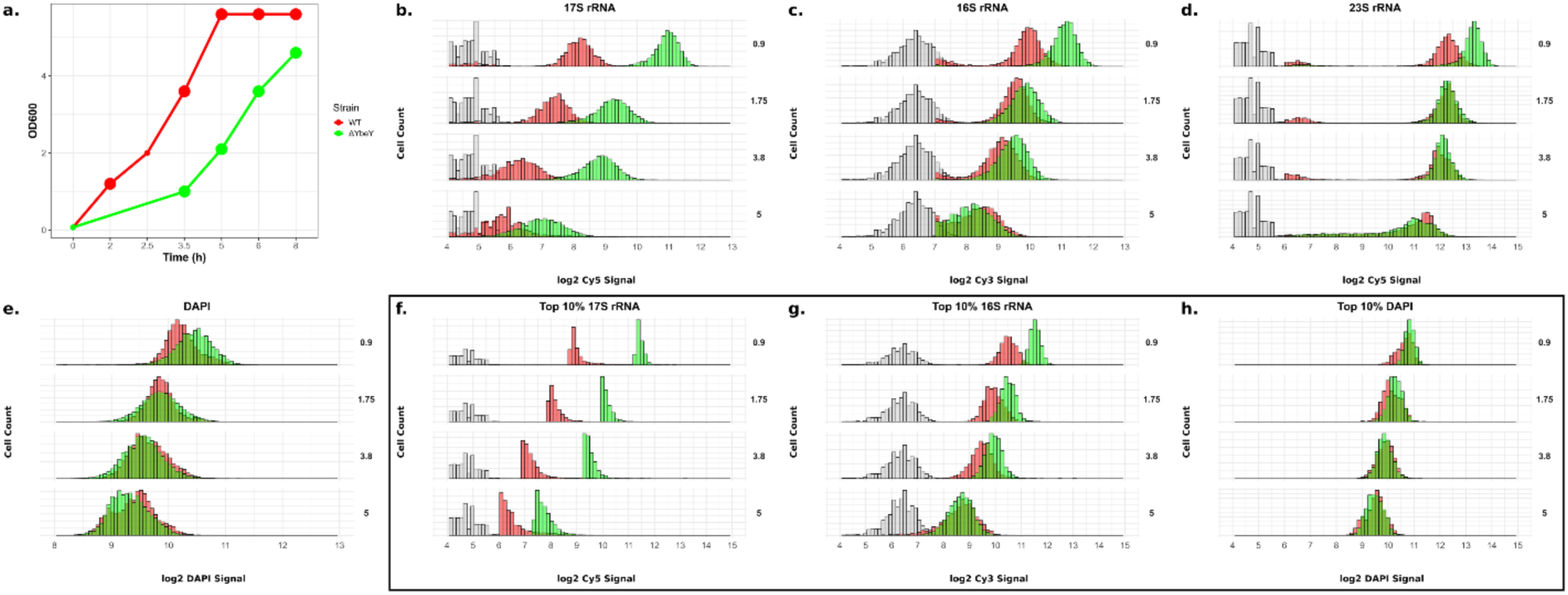
Flow cytometry reveals rRNA levels at single-cell resolution. a. Growth curves for the BW25113 (WT; in red) and ΔybeY (in green) strains at OD_600_ nm. The large dots show sampling times (as depicted in panels b.-h.). b-d. Distributions of 17S pre-rRNA, 16S rRNA and 23S rRNA levels at single-cell resolution, shown at time points indicated on panel a. Non-stained control refers to WT cells treated with a mock solution instead of fluorescent probes. The fluorescence signal corresponding to RNA levels is displayed on a log2 scale. e. DAPI signal (DNA levels), f.-g. The signal from cells in the top 10% according to 17S rRNA levels. f. shows 17S rRNA signal and g. shows 16S rRNA signal from the same cell population h. DAPI signal at the top 10% population (from the same cells as in f and g).

WT cultures displayed low 17S signal throughout, similar to their behavior in MOPS, with a further decline into the stationary phase. In contrast, *ΔybeY* cells displayed a markedly higher 17S signal during early growth — on average approximately 8-fold above WT — and a narrow, well-defined unimodal distribution, indicating uniform 17S pre-rRNA accumulation across cells. 17S signal in *ΔybeY* declined gradually in the stationary phase, reaching WT levels by the end of the time course (Figure 5b). This uniform, early-growth 17S pre-rRNA accumulation is qualitatively different from the late-phase, highly heterogeneous 17S accumulation seen in *ΔybeX* under Mg²⁺ limitation (Figure 1b): nearly all *ΔybeY* cells accumulate 17S pre-rRNA simultaneously, whereas in *ΔybeX* the 17S accumulation is confined to a variable subpopulation and the overall distribution broadens by up to ∼25-fold.

As in the *ΔybeX* analysis, we examined the top 10% of cells by 17S pre-rRNA signal. *ΔybeY* cells with the highest 17S pre-rRNA content also carried the highest 16S rRNA content. As in the full population, the top 10% subpopulation showed a decline in signal as cultures approached the stationary phase (Figure 5f,g). DAPI signal distributions for the full population (Figure 5e) and top-10% subpopulation (Figure 5h) are also shown.

Aggregate analysis across replicates confirmed that during early growth, *ΔybeY* carried more of every rRNA species — approximately 2-fold more 16S and 23S rRNAs, and approximately 8-fold more 17S pre-rRNA — than WT. By the late growth phase, the two strains carried comparable rRNA levels (Supplementary Figure 13). Correlations for the top 10% subpopulation are shown in Supplementary Figure 14.

Plotting the single-cell-level 16S/23S and 17S/16S rRNA ratios sharpened the contrast between the two mutants. As in *ΔybeX*, the 23S/16S ratio distribution in *ΔybeY* was tightly balanced, with a median ratio of 1.8–2.2 and 10th–90th-percentile fold-range of 1.3–1.6 in both strains, suggesting tight stoichiometries in mature rRNAs (Figure 6). The 17S/16S rRNA ratio distribution, however, was uniformly elevated in *ΔybeY* by ∼2.8-fold over WT across exponential growth (representative medians 1.15 vs 0.40, 0.87 vs 0.32 and 0.69 vs 0.24 at OD₆₀₀ < 1, 1–2 and 3–4, respectively), declining toward the WT value by the onset of stationary phase (0.37 vs 0.21 at OD₆₀₀ ≥ 4). Crucially, and unlike *ΔybeX*, this elevation reflected a coherent shift of the whole population: the *ΔybeY* 17S/16S distribution stayed narrow and unimodal (fold-range ≈ 1.5–2.7, comparable to WT and far below the ≈ 8.5-fold spread of late-exponential *ΔybeX*). The single-cell ratio distribution therefore clearly separates the two phenotypes — a uniform, population-wide accumulation of pre-rRNA in *ΔybeY* versus a subpopulation-restricted, highly heterogeneous accumulation in *ΔybeX* (Figure 6f; individual experiments and the pooled composite in Supplementary Figures 11, 12).

**Figure 6.**
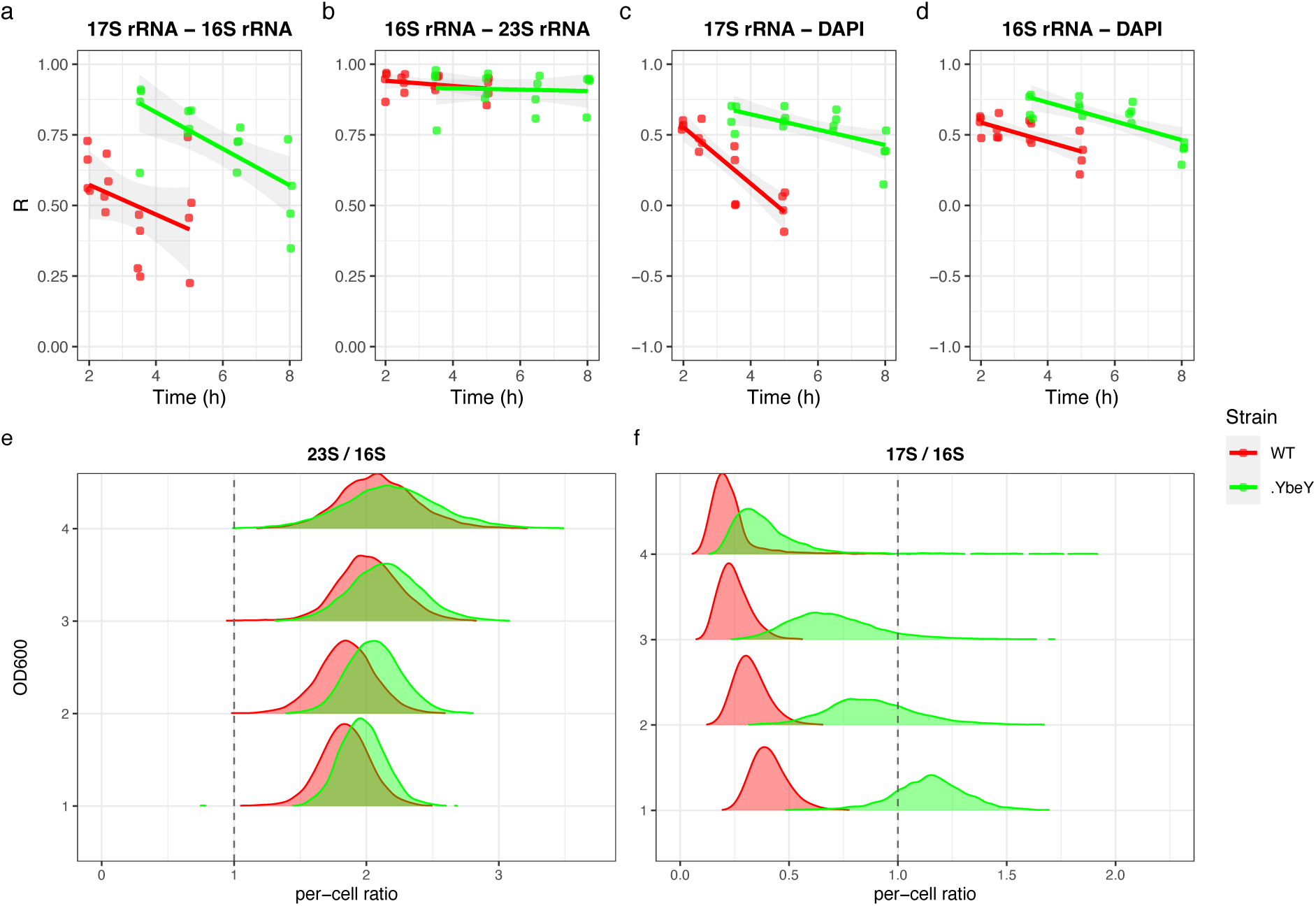
The stoichiometry of a) 17S vs. 16S rRNAs, b) 16S vs. 23S rRNAs, c) 17S vs DAPI, and d) 16S vs DAPI from full data, assessed by linear correlation. For each time point and strain in each experiment, Pearson correlation coefficients (R) were calculated, and temporal dynamics were modeled by pooled linear regression. Data points from independent experiments are represented by points (red for WT and green for ΔybeY) and regression lines with shaded 95% CI are colored accordingly. e) Per-cell rRNA-ratio distributions (linear scale; dashed line = ratio of 1) for 23S/16S and f) 17S/16S ratios for WT (red) and ΔybeY (green) across OD₆₀₀ bins. Individual experiments and the pooled composite are in Supplementary Figures 11 and 12.

## DISCUSSION

Using rRNA fluorescence *in situ* hybridization coupled with flow cytometry (rRNA-FISH-flow), we have quantified 16S rRNA, 23S rRNA, and 17S pre-rRNA at single-cell resolution in *E. coli* BW25113 and two isogenic deletion mutants of genes encoded in the *ybeZYX-lnt* operon: *ΔybeX* and *ΔybeY*. Four principal findings emerge. First, *ΔybeX* cells grown into the late exponential phase under Mg^2+^ limitation develop striking cell-to-cell heterogeneity in 17S pre-rRNA, with approximately 25-fold differences between the lowest- and highest-signal cells. Second, upon regrowth from the stationary phase, *ΔybeX* cultures display a distinctly bimodal 17S distribution throughout the extended lag phase, with the population gradually shifting from a high-17S to a low-17S mode as visible growth resumes. Third, *ΔybeY* cultures grown in rich medium accumulate 17S pre-rRNA during early exponential growth in a narrow, unimodal distribution that is ∼8-fold above wild type and is progressively cleared as cells enter stationary phase. Fourth, the stoichiometry between mature 23S and 16S rRNAs is tightly maintained in both mutants throughout all conditions tested, indicating that the defect lies at the level of pre-rRNA processing and synthesis. Together, these data demonstrate that YbeX and YbeY perturb ribosome biogenesis through qualitatively different routes, and that the extended lag phase of *ΔybeX* reflects a subpopulation-specific, rather than uniform, clearance of pre-ribosomal intermediates.

### Single-cell analysis reveals subpopulation structure

The bulk-level phenotypes of *ΔybeX* and *ΔybeY* — accumulation of 17S pre-rRNA and of 16S rRNA degradation intermediates, extended lag phase in *ΔybeX* (Sarıgül et al. 2024), and constitutive 17S pre-rRNA accumulation in *ΔybeY* (Davies et al. 2010; Jacob et al. 2013)— have been described in some detail. However, these studies cannot distinguish whether an observed RNA species reflects a modest uniform per-cell defect or a severe defect confined to a subpopulation. Our single-cell measurements help to resolve this ambiguity. In *ΔybeX*, the mean rise in 17S pre-rRNA levels conceals a broadening of the 17S pre-rRNA distribution from ∼5-fold to ∼25-fold cell-to-cell variation. Moreover, during outgrowth, the mean 17S pre-rRNA trajectory masks a switch-like transition between two coexisting subpopulations.

This observation helps to explain the colony-size heterogeneity reported by Sarıgül et al. (2024) when *ΔybeX* stationary cells were plated on LB agar. We believe that colony-size heterogeneity on plates and 17S-signal bimodality in liquid medium outgrowth are two faces of the same process, whereby within a genetically homogeneous *ΔybeX* stationary culture, individual cells inherit qualitatively different pre-ribosomal states, which in turn lead to divergent regrowth kinetics.

### *ΔybeX* and *ΔybeY* act on ribosome biogenesis through distinct pathways

The most informative contrast in our data is not between mutant and wild type, but between *ΔybeX* and *ΔybeY*. Despite being co-encoded in the *ybeZYX-lnt* operon and functionally converging on the ribosome, the two deletion mutants produce qualitatively different single-cell 17S pre-rRNA signatures. *ΔybeY* shows uniform, constitutive, medium-independent accumulation of 17S pre-rRNA across nearly all cells during early exponential growth, with a narrow unimodal distribution, while *ΔybeX* shows a late-phase, Mg^2+^-dependent accumulation that is confined to a variable subpopulation and is accompanied by a broadening of the distribution. These signatures are consistent with two distinct modes of action.

For YbeY, a direct enzymatic role in 16S 3ʹ-end maturation has been proposed (Davies et al. 2010; Jacob et al. 2013), in which every cell that transcribes pre-rRNA experiences a similar defect. The observed tight unimodal distribution of 17S pre-rRNA levels is expected under such a direct biochemical mechanism. By contrast, for YbeX, the defect manifests only under Mg^2+^ limitation, hypothetically after substantial cytoplasmic Mg^2+^ has been depleted, and only in a subset of cells. This is expected of an indirect defect whose penetrance depends on each cell’s physiological state. The YbeX membership of the CorC family of membrane-proximal Mg^2+^/divalent-cation homeostasis factors is consistent with this interpretation (Huang et al. 2021; Iwadate and Slauch 2025), as is the alleviation of its ribosomal phenotype by Mg^2+^ supplementation (Sarıgül et al. 2024). The most parsimonious model is that YbeX contributes to intracellular Mg^2+^ homeostasis and that the ribosomal phenotype is a downstream consequence of Mg^2+^ insufficiency at late growth phases, when Mg^2+^ demand outpaces supply.

This suggests a more general principle: tight and heterogeneous cell-to-cell distributions can serve as flags for distinct mechanisms of action. Direct biochemical lesions tend to produce tight distributions because every cell is equally affected; indirect lesions that operate through variable upstream cellular states (intracellular Mg^2+^, (p)ppGpp levels, membrane potential) tend to produce heterogeneous distributions because the upstream state differs between sister cells.

### The defect in *ΔybeX* cells is post-transcriptional and temporally decoupled from growth arrest

Our work is consistent with a role of YbeX at a post-transcriptional (processing/clearance) step, while the elevated mature-rRNA content and the failure to down-regulate rRNA at the transition indicate an accompanying failure to restrain rRNA output — both plausibly downstream of dysregulated Mg^2+^ homeostasis. In both strains, the cell-to-cell correlation between mature 23S and 16S rRNAs remains high throughout growth, consistent with balanced transcription of rRNA from the *rrn* operons. By contrast, the 17S–16S rRNA correlation in *ΔybeX* varies with growth phase and breaks down in the outgrowth experiment at the point where accumulated 17S pre-rRNA is being cleared. We conjecture that new rRNA transcription remains coupled across both subunits in every cell, but that the fate of individual 17S pre-rRNA molecules, processed or retained, diverges between cells.

The 17S pre-rRNA accumulation in *ΔybeX* is temporally decoupled from growth arrest. 17S pre-rRNA levels already diverge between *ΔybeX* and wild type in mid-exponential phase, but the cultures continue to grow at indistinguishable rates for several more hours (Figure 1a, b, f). Only at the late-exponential/early stationary phase transition does *ΔybeX* fail to down-regulate mature rRNA and goes into growth arrest. This lag between 17S pre-rRNA accumulation and growth failure suggests that 17S pre-rRNA accumulation is not, *per se*, immediately translationally limiting. Rather, growth arrest appears to require the crossing of a threshold — potentially in the ratio of mature to immature subunits, in the depletion of a Mg^2+^-dependent cofactor pool, or in the accumulated load of stalled pre-ribosomes.

### Origin of heterogeneity and the bimodal outgrowth

Cell-to-cell heterogeneity in rRNA levels is not unique to our mutants: all strains, including wild type, exhibit roughly 10- to 15-fold variation in mature 16S and 23S rRNA signals at every time point examined. A substantial literature documents the sources of such baseline variability, including stochastic gene expression (Uphoff et al. 2016), variation in gene and operon copy number between cells (Kunnimalaiyaan et al. 2001; Wang et al. 2005), variable activity of individual *rrn* operons (Anda et al. 2023), heterogeneity in ribosome composition (Natchiar et al. 2017; Shen et al. 2025) and noise in downstream translation (Golding et al. 2005).

A recent study in *Staphylococcus aureus* shows lower 16S rRNA levels and lower cell to cell variation in stationary phase than in exponential phase rRNA (Tripathi et al. 2025). Our baseline measurements are consistent with this work. What distinguishes *ΔybeX* is a switch-like transition during outgrowth. Unlike the continuous broadening seen in growth round-one experiments, the round-two (regrowth experiment) 17S pre-rRNA distribution in *ΔybeX* is distinctly bimodal: two peaks appear at distinct timepoints, and the relative sizes of the peaks shift over time, with a tendency toward low 17S pre-rRNA signal once growth resumes after the extended lag phase. This behavior can be explained by phenotypic bistability, characterized by two underlying cellular states that are individually stable on the timescale of sampling, with cells transiting from the high-17S to the low-17S state as the lag phase progresses. The point at which visible growth resumes coincides approximately with the point at which the two subpopulations are equally populated (5-6h), suggesting that the population-level lag reflects the time required for half of the cells to transit from the high-17S to the low-17S state.

This interpretation places the *ΔybeX* lag phenotype in the broader context of bacterial bet-hedging and persistence. Slow-growing or dormant subpopulations that tolerate antibiotic exposure better than their faster-growing sisters are a recurring feature of bacterial populations and are frequently accompanied by reduced ribosome content or altered translation state (Balaban et al. 2004; Maisonneuve and Gerdes 2014). Sarıgül et al. (2024) reported that *ΔybeX* cells are more sensitive to ribosome-targeting antibiotics than wild type — a phenotype that can be rationalized if the high-17S pre-rRNA subpopulation carries a disproportionate pool of immature, antibiotic-binding pre-ribosomes. A testable prediction is therefore that the high-17S subpopulation coincides with the cells that display the antibiotic and temperature sensitivities. Direct tests would require sorting cells on 17S pre-rRNA signal and retesting antibiotic sensitivity and regrowth kinetics in the sorted fractions.

### Future directions

While our single-cell approach to steady-state ribosomal RNA quantification is useful for bringing cell-to-cell heterogeneity in ribosomal metabolism into focus, its main shortcoming is that it fails to provide a dynamic picture of ribosomal homeostasis. We cannot, therefore, differentiate between plausible hypotheses of the YbeX protein function: whether it’s a Mg-sensing protein that is directly involved in regulating rRNA operon transcription, or is its main function in driving ribosomal rRNA processing and/or ribosomal assembly under low Mg^2+^ conditions. To resolve these questions, kinetic experiments on rRNA transcription and 17S-to-16S rRNA processing, as well as on rRNA operon transcription and its Mg-dependent regulation, and direct measurements of Mg^2+^ at the single-cell level are needed. We believe that the molecular function of the YbeX protein in ribosomal metabolism is both a promising and a worthwhile avenue for further study.

## MATERIALS AND METHODS

### Growth media

MOPS minimal medium (Neidhardt et al. 1974), supplemented with 0.4% glucose and 10 mM or 50 µM MgCl_2_, was used for growing WT and *ΔybeX* strains, and the medium was filter-sterilized through a 0.22 µm filter. LB (BD Difco^TM^ LB Broth #240230) medium was used to culture WT and *ΔybeY* strains; autoclaving was performed at 121°C for 15 min.

### Strains used in the study

*Escherichia coli* wild-type strain BW25113 and the isogenic mutants *ΔybeX* and *ΔybeY* were used. BW25113 and *ΔybeY* were obtained from the Keio collection (Baba et al. 2006), *ΔybeX* was constructed previously (Sarıgül et al. 2024). *ΔybeX* and *ΔybeY* carry a kanamycin resistance gene cassette as a selective marker; accordingly, all mutant strains’ cultures were supplemented with kanamycin at a final concentration of 50 µg/mL. All strains were streaked from glycerol stocks onto LB agar plates (BD Difco^TM^ LB Agar #240110) and incubated overnight at 37°C.

### Preparation and growth of cells for FISH-FLOW

#### Growth under Mg^2+^ limitation

A single colony was inoculated into 10 mL MOPS minimal medium supplemented with 0.4% glucose and 10 mM MgCl_2_. Cells were grown overnight at 37°C with orbital shaking at 200 rpm to support optimal growth prior to Mg^2+^ limitation. From each overnight culture, 1 mL was transferred into 1.5 mL microcentrifuge tube. Cells were pelleted at 5,000 × g for 5 minutes at room temperature and the pellet was resuspended in 1 mL MOPS minimal medium lacking MgCl_2_. Cells were re-pelleted under the same centrifugation conditions, and this washing step was repeated for a total of three washes, to remove residual Mg^2+^ from the pre-culture medium.

Following the final wash, cells were resuspended in 50 mL MOPS minimal medium supplemented with 0.4% (w/v) glucose and 50 µM MgCl_2._ Cultures were initiated at starting OD_600_ of 0.05-0.06 in 50 mL of medium and grown at 37°C with orbital shaking at 200 rpm. All experiments were performed in at least three biological replicates.

### Stationary phase re-inoculation

Cells were collected from the early stationary phase to restart growth in MOPS supplemented with 50 µM MgCl_2_. 1 mL of culture was centrifuged at 5,000 × g at room temperature for 5 min. Pellets were resuspended in fresh growth medium. Fresh culture was initiated in 70 mL medium at an OD_600_ of 0.1. Cultures were grown at 37°C with orbital shaking at 200 rpm up to 9 hours.

#### Growth in LB Medium

A single colony was inoculated into 10 mL of LB medium. Cells were grown overnight at 37°C, and a 1 mL sample was pelleted at 5,000 × g, room temperature for 5 minutes. Cells were resuspended in 1 mL LB and inoculated into 50 mL fresh LB medium at a starting OD_600_ of 0.05-0.06. Cultures were grown at 37°C until the stationary phase.

Samples for FISH analysis were collected at the time points indicated in Figures 1, 3, and 5.

### FISH-FLOW

We adapted the protocol of Mattioli and Avraham (2024) with minor changes.

#### Fixation and Permeabilization

At each time point, 1 mL of culture was centrifuged at 5,000 × g at room temperature for 5 minutes. The pellet was resuspended once in 1 x PBS and centrifuged under the same conditions. Cells were resuspended in fixation buffer (1 x PBS in 4% formaldehyde; Table 1). For fixation, cells were incubated at 4°C for 30 minutes with gentle shaking on a nutator. Fixed cells were washed twice with 1 mL 1 x PBS, pelleting at 5,000 x g for 5 minutes at 4°C between washes. Post-fixation centrifugation was performed at 4°C to preserve cellular integrity.

Cell pellets were resuspended in 300 µL ddH_2_O. Ethanol was added to a final concentration of 70% for permeabilization. Cells were incubated overnight at 4°C with gentle shaking on a nutator.

#### Probe Hybridization

Following permeabilization, cells were pelleted at 5,000 x g for 10 minutes at RT. Supernatant was discarded, and cells were resuspended in wash buffer (2 x SSC, 45% Formamide (AppliChem)) and incubated at room temperature for 15 minutes with gentle agitation on a nutator. Cells were re-pelleted and resuspended in 50 µL hybridization buffer composed of 10% dextran sulfate (Sigma), 2 x SSC, 45% Formamide, 0.02% BSA, 100 µg/mL yeast tRNA, supplemented with 0.04 U/µL RiboLock RNase inhibitor (Thermo Fisher). Fluorescently labeled probes were added to the hybridization mixture at a final concentration of 1 µM per probe. Cells were incubated at 37°C overnight to allow hybridization to target rRNA sequences.

Three probe sets were used in this study (23S-16S/17S-16S/Control). To increase the sensitivity of fluorescent signal detection, two separate probes binding to distinct sites were designed against each rRNA species (23S and 16S rRNA) and against 17S pre-rRNA. To assess non-specific background hybridization, a control probe set was designed with no complementarity to any *E. coli* genomic sequence. Probes sequences and fluorophore conjugations are listed in Table 2.

#### Preparation of Samples for FACS Analysis

After hybridization, cells were resuspended in 1 x PBS and pelleted by centrifugation at *5,000 x g*, for 10 minutes at room temperature. The supernatant was aspirated to approximately 100 µL, and cells were resuspended in 1 x PBS and re-pelleted. The pellets were resuspended in 1 x PBS. For DAPI staining, cells were diluted 1:10 in 1× PBS containing DAPI (Invitrogen) to a final concentration of 10 µg/mL. Cells were incubated at room temperature for 30 minutes to allow DAPI staining. DAPI staining was performed in the dark. Prior to acquisition, additional dilutions were done as necessary based on event rate to achieve 10^5^-10^6^ cells per analysis.

##### FACS Analysis

Flow cytometry was performed on a BD LSR Fortessa instrument. A forward-scatter (FSC) threshold of 700 was applied to reduce acquisition of low-intensity background events. Fluorescence was measured using the red laser for Cy5, the yellow–green laser for Cy3, and the violet laser for DAPI (Table 3).

Data were analyzed in R version 4.3. Single cells were identified based on FSC-A versus FSC-H gating (Supplementary Figure 15). Downstream analyses were restricted to the dense singlet population (10^3^ ≤ FSC-A, FSC-H ≤ 10^3.5^). Events with low FSC values, which exhibited poor FSC-A/FSC-H correlation and were consistent with debris/noise, as well as high-FSC events corresponding to large cells, filaments, or aggregates, were excluded. No compensation was applied because the spectral overlap among the Cy3, Cy5, and DAPI channels was negligible, given their spectral separation.

Plots and statistical analyses were generated using the ggplot2 package.

##### Estimation of Growth Rates

Growth rates and lag times were estimated from OD_600_ growth curves by linear regression of ln(OD_600_) versus time during the exponential phase. The specific growth rate (µ) was taken as the slope of the regression, and the lag time was defined as the time at which the fitted line intersected the initial OD_600_ value. Analyses were performed in R (version 4.3).

## Data and code availability

Generative AI tools (Claude Opus 4.7 and Claude Opus 4.8, Anthropic) were used to assist with R code development for growth curve analysis and figure preparation. All AI-generated content was reviewed, verified, and accepted by the corresponding author, who takes full responsibility for the manuscript content. Data and code to reproduce the figures are available at the datadoi.ee (DOI: https://doi.org/10.23673/UKLCFH).

## ACKNOWLEDGEMENTS

We would like to thank Marilis Hinnu for her comments on flow cytometry, Aili Togoma and Brita Laht for their technical support and help in using and adjusting BD LSRFortessa flow cytometer, and Niilo Kaldalu for helpful discussions. This work was supported by the Estonian Research Council grant PRG2696.

## AUTHOR CONTRIBUTIONS

Abbas Mansour developed a methodology and conducted experiments, analyzed and interpreted the data, and wrote the original draft. İsmail Sarıgül: study design, data interpretation, writing. Tanel Tenson: study design, writing. Ülo Maiväli: study design and supervision, writing.

## SUPPLEMENTARY FIGURES

**Supplementary Figure 1.**
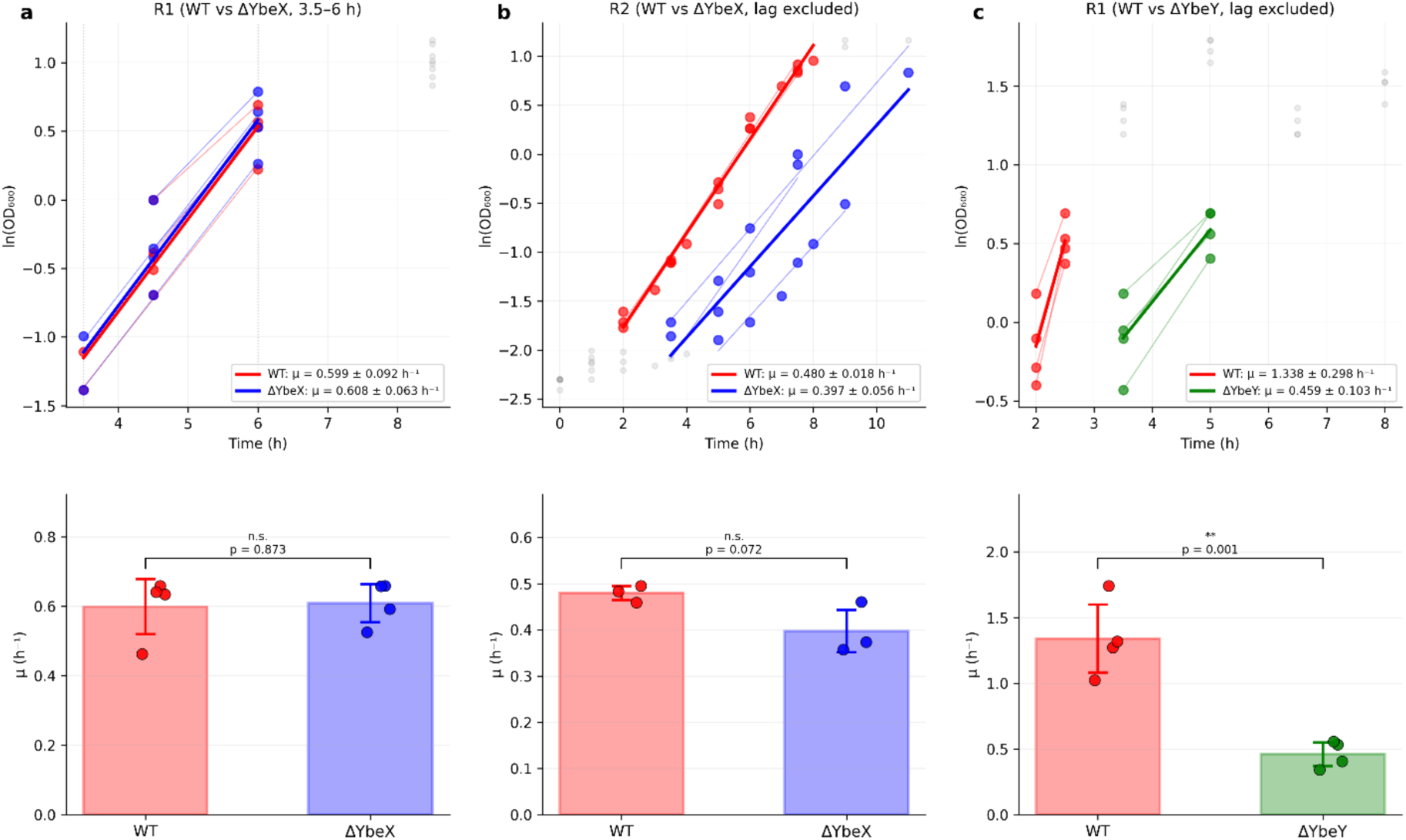
Exponential growth of BW25113, ΔybeX and ΔybeY strains. a) Linear regression of ln(OD_600_) against time of wt and ΔybeX in the first growth round. The bar plots show exponential growth rates per hour (µ,h^-1^)). Both strains show similar growth rates (p = 0.87, Student’s t-test). b) Linear regression of wt and ΔybeX in the second round of growth. No statistically significant difference in the exponential growth rate was observed between the two strains (p = 0.07). c) Linear regression of wt and ΔybeY, with growth rates shown as bar plots, ΔybeY grew ∼3-fold slower than wt (p = 0.001). Data points represent individual biological replicates; error bars indicate SD. Lag-phase and stationary-phase data points were excluded from the regression fits.

**Supplementary Figure 2.**
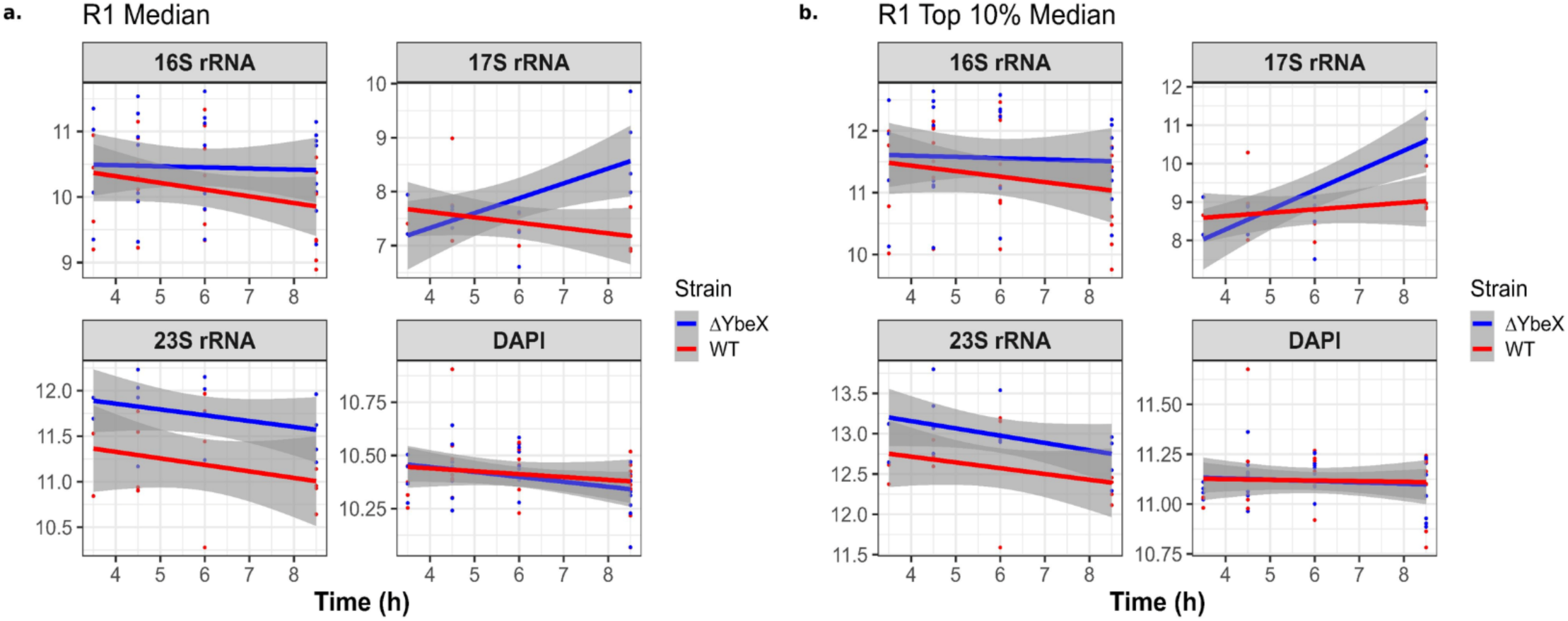
Aggregate-level analysis of all flow cytometry experiments. a. Linear regression of median rRNA and DAPI signals. The median rRNA levels for each independent experiment are plotted against growth time from the start of the experiment (dilution of overnight culture into Mg-limited media). b. Linear regression of interquartile ranges (IQR) of each sample. Data points from independent experiments are represented by points (red for WT and blue for ΔybeX) and regression lines with shaded 95% CI are colored accordingly.

**Supplementary Figure 3.**
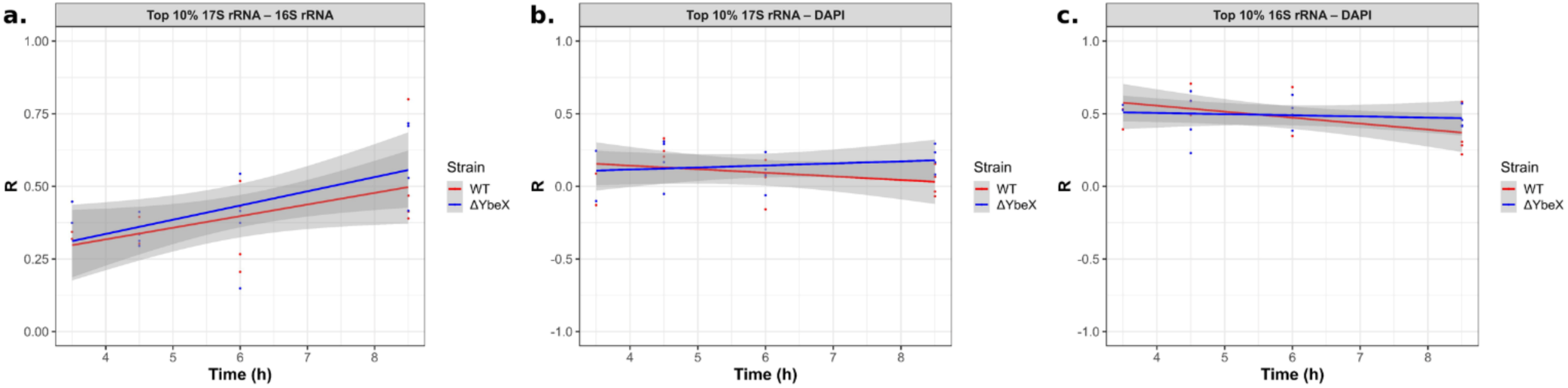
Top 10% (by 17S signal) correlation between a) 17S-16S, b) 17S-DAPI, c) 16S-DAPI. Data points from independent experiments are represented by points (red for WT and blue for ΔybeX) and regression lines with shaded 95% CI are colored accordingly.

**Supplementary Figure 4.**
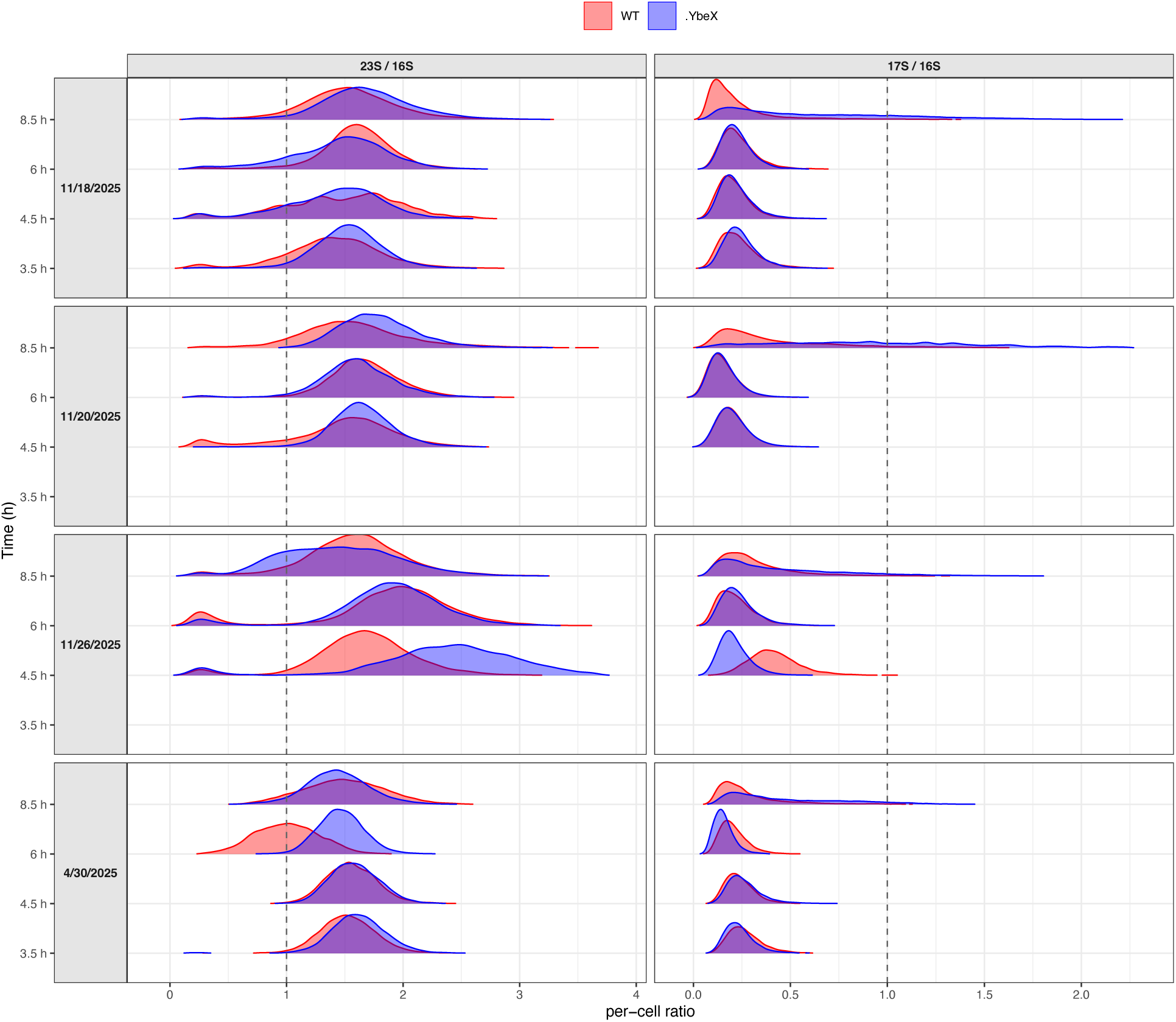
Per-cell 23S/16S and 17S/16S rRNA ratios in WT and ΔybeX, first growth cycle (Mg^2+^-limited MOPS). Linear-scale ridgelines of the per-cell ratio (dashed line = ratio of 1); WT in red, ΔybeX in blue. Each independent experiment in its own panel (faceted by date), time on the y-axis. Note the wide distribution of the ΔybeX 17S/16S at the late time point in every replicate, compared to the narrow 23S/16S. The dates of experiments are shown on the second Y-axis and correspond with those in the supplementary data.

**Supplementary Figure 5.**
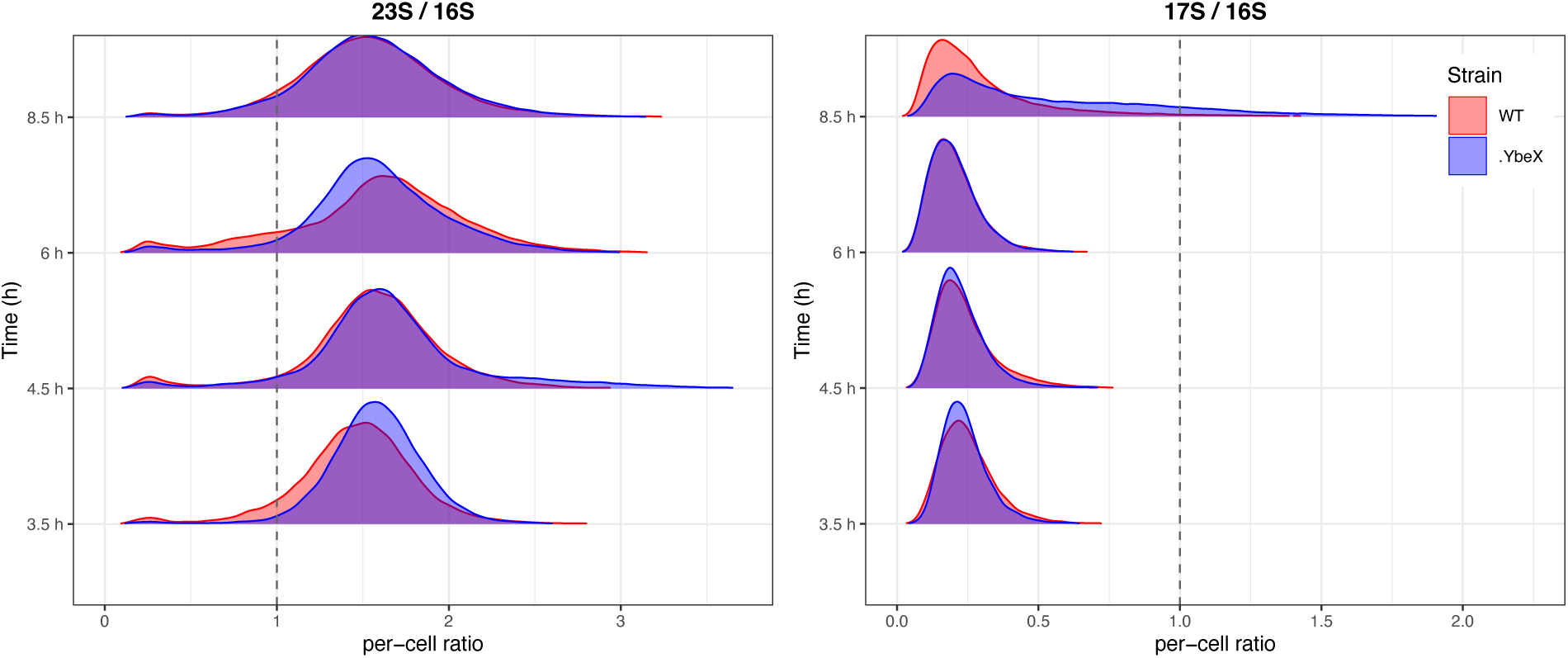
All replicates pooled within each time point. Note the marked broadening of the ΔybeX 17S/16S distribution at the late time point, against the stable, narrow 23S/16S ratio.

**Supplementary Figure 6.**
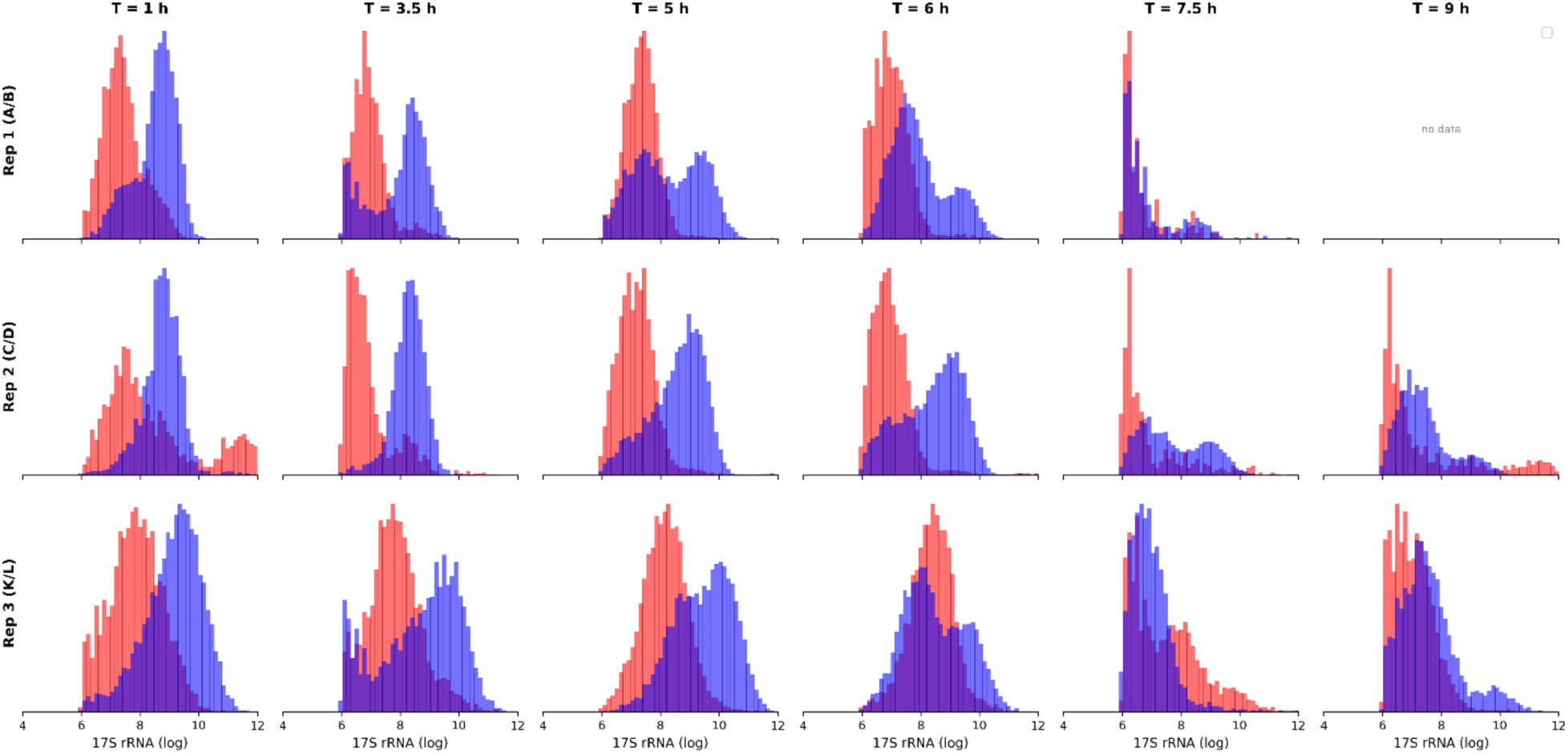
Distribution of 17S rRNA in both BW25113 (red) and ΔybeX (blue) from the second growth round experiment. Three independent experiments are shown. T = 1 h indicates the one hour time point, and so on. The rRNA species is indicated on the X-axis label. All values are in log2 units.

**Supplementary Figure 7.**
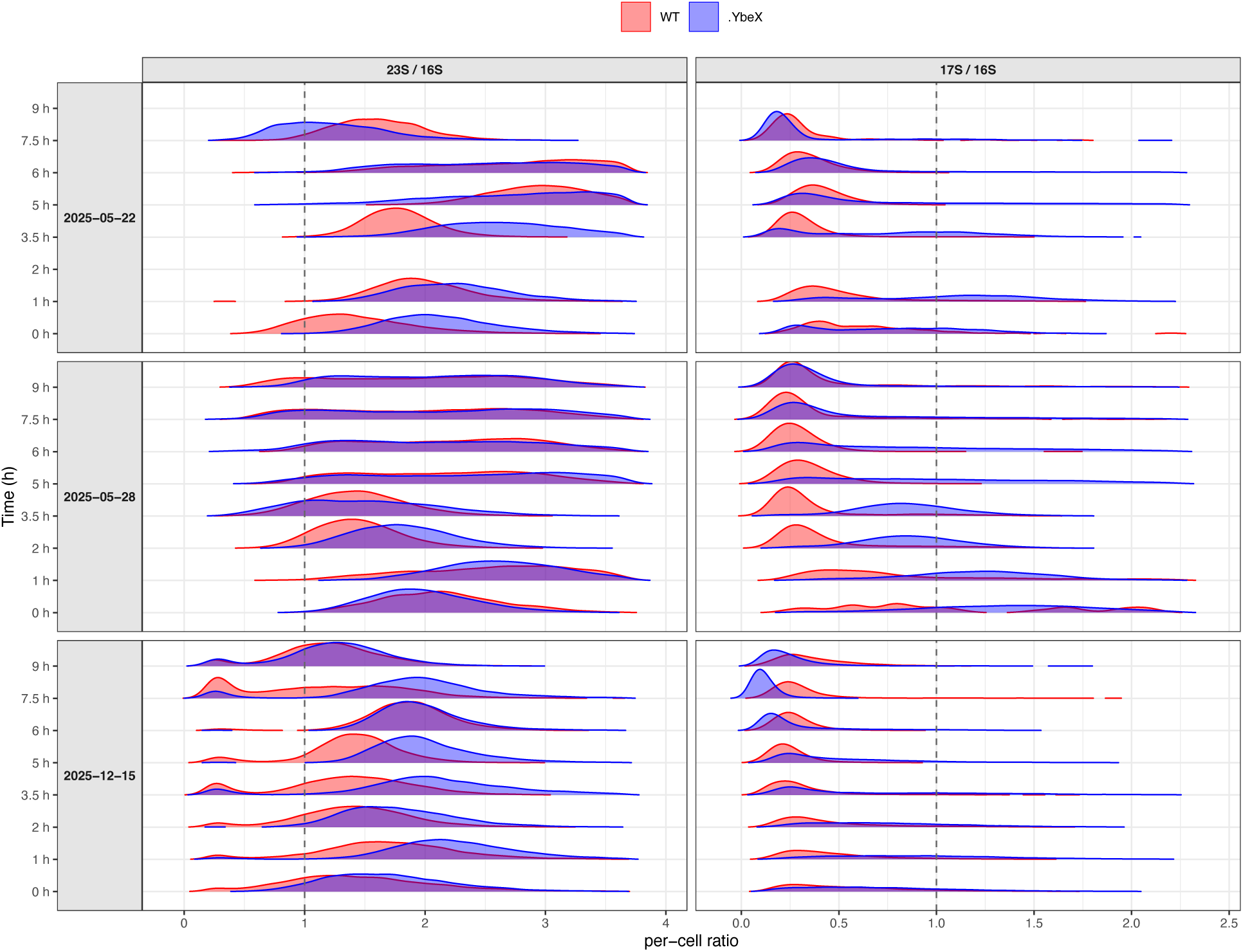
Distributions of per-cell 23S/16S and 17S/16S rRNA ratios in WT and ΔybeX, second growth cycle (outgrowth from stationary phase) The ΔybeX 17S/16S ratio is high and broadly distributed during the lag phase and collapses toward the WT value as visible growth resumes; the 23S/16S ratio stays narrow throughout. The second Y-axis indicates the experiment date, which corresponds to the date in the supplementary data table.

**Supplementary Figure 8.**
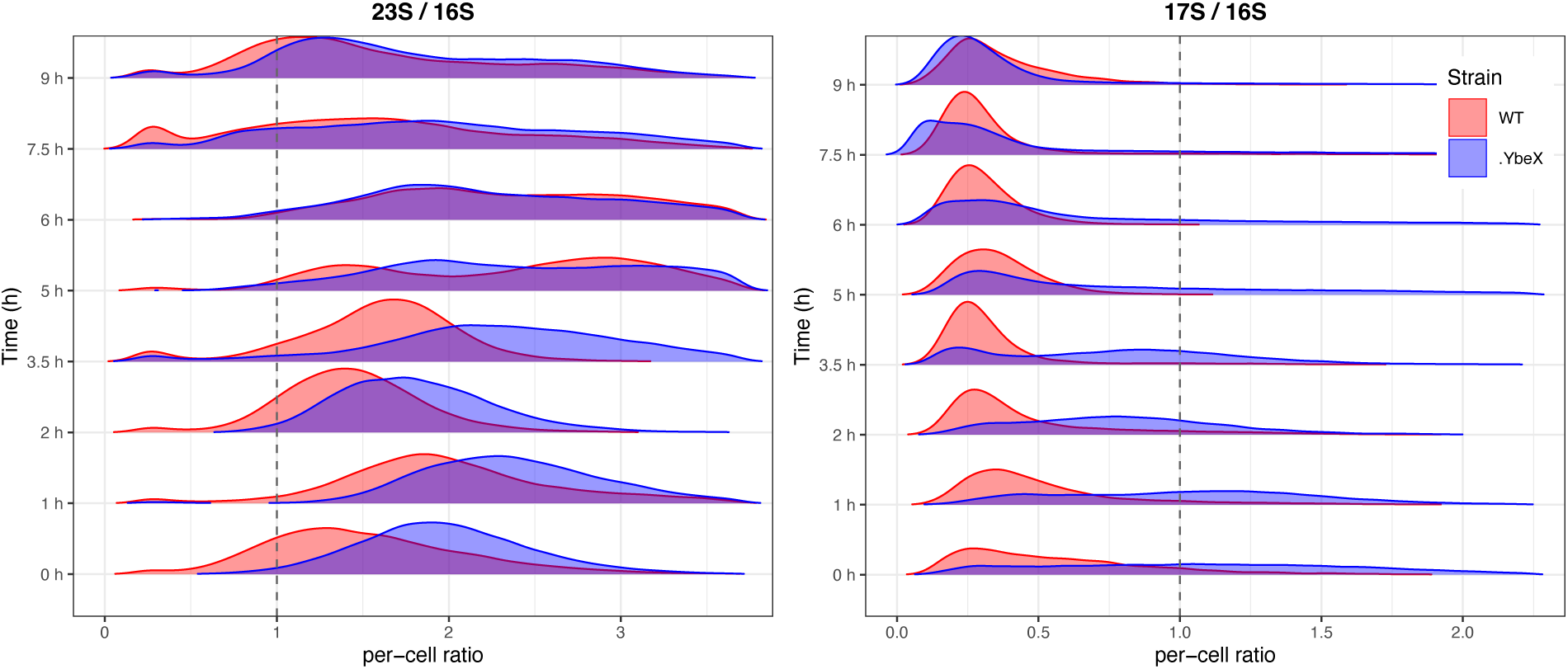
Distribution of per-cell ratios of 23S/16S and 17S/16S rRNAs from the pooled data of the second-round growth experiment.

**Supplementary Figure 9.**
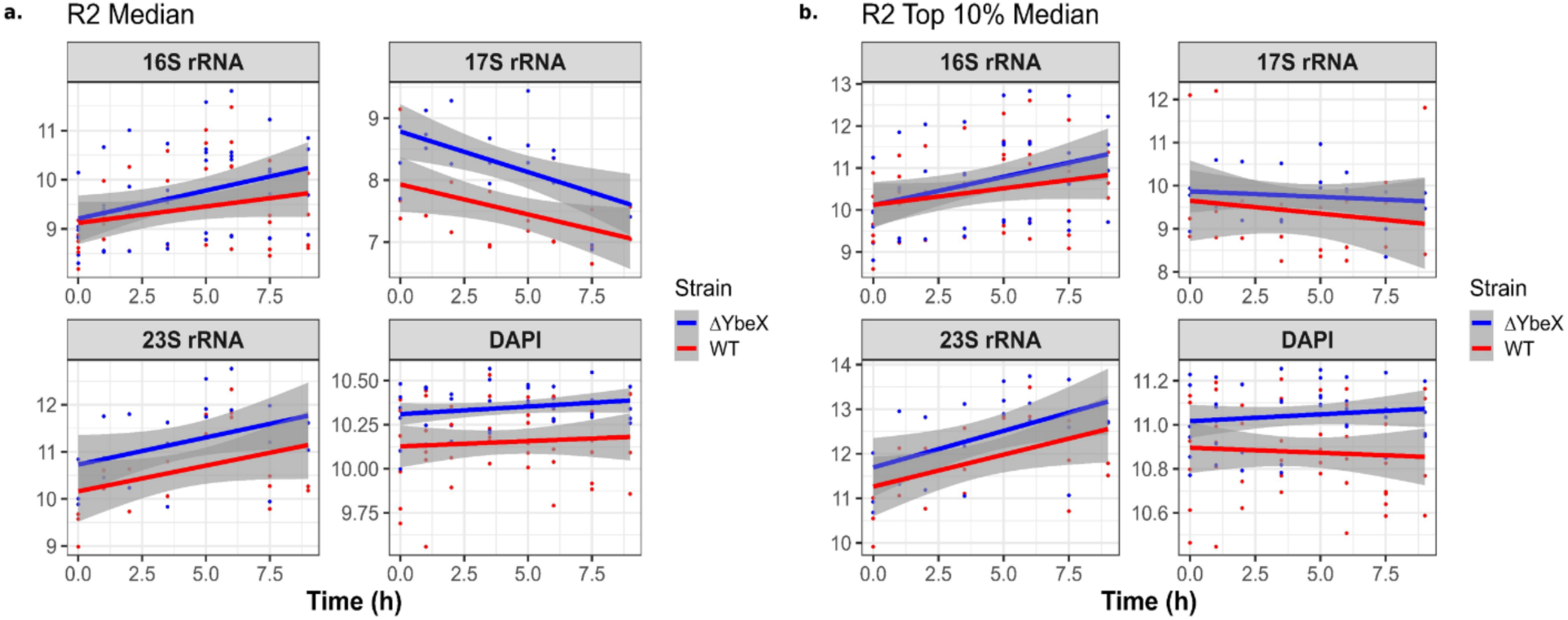
Aggregate-level analysis of the temporal evolution of median rRNA levels and variation in the second growth cycle experiment. Data points from independent experiments are represented by points (red for WT and blue for ΔybeX) and regression lines with shaded 95% CI are colored accordingly. accordingly.

**Supplementary Figure 10.**
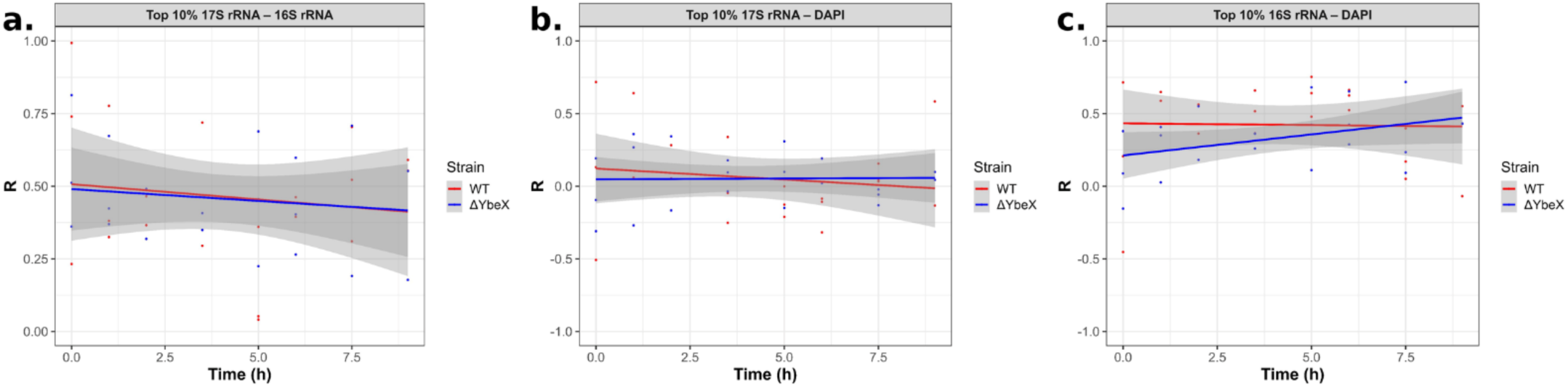
Linear regression model for correlation between WT and ΔybeX cells manifesting top 10% signal from the second growth culture: a) 17S vs 16S, b) 17S vs DAPI, c) 16S vs DAPI.

**Supplementary Figure 11.**
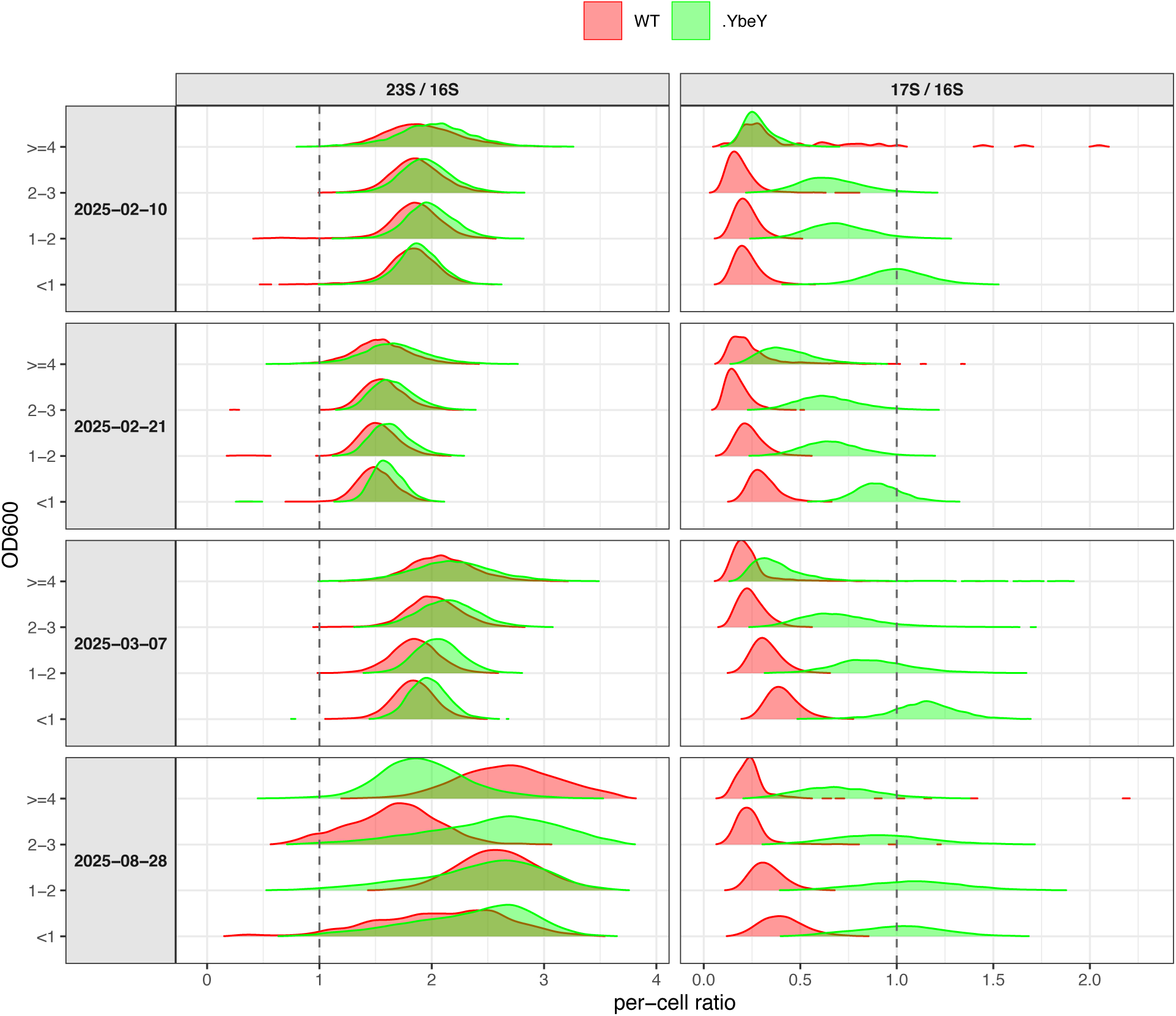
Distribution of per-cell 23S/16S and 17S/16S rRNA ratios in WT and ΔybeY strains, at indicated OD_600_ values. Layout is as in Supplementary Figure 4 with ΔybeY shown in green and OD_600_ on the y-axis. The ΔybeY 17S/16S ratio is uniformly elevated as a narrow, unimodal distribution, in contrast to the heterogeneous broadening seen in ΔybeX.

**Supplementary Figure 12.**
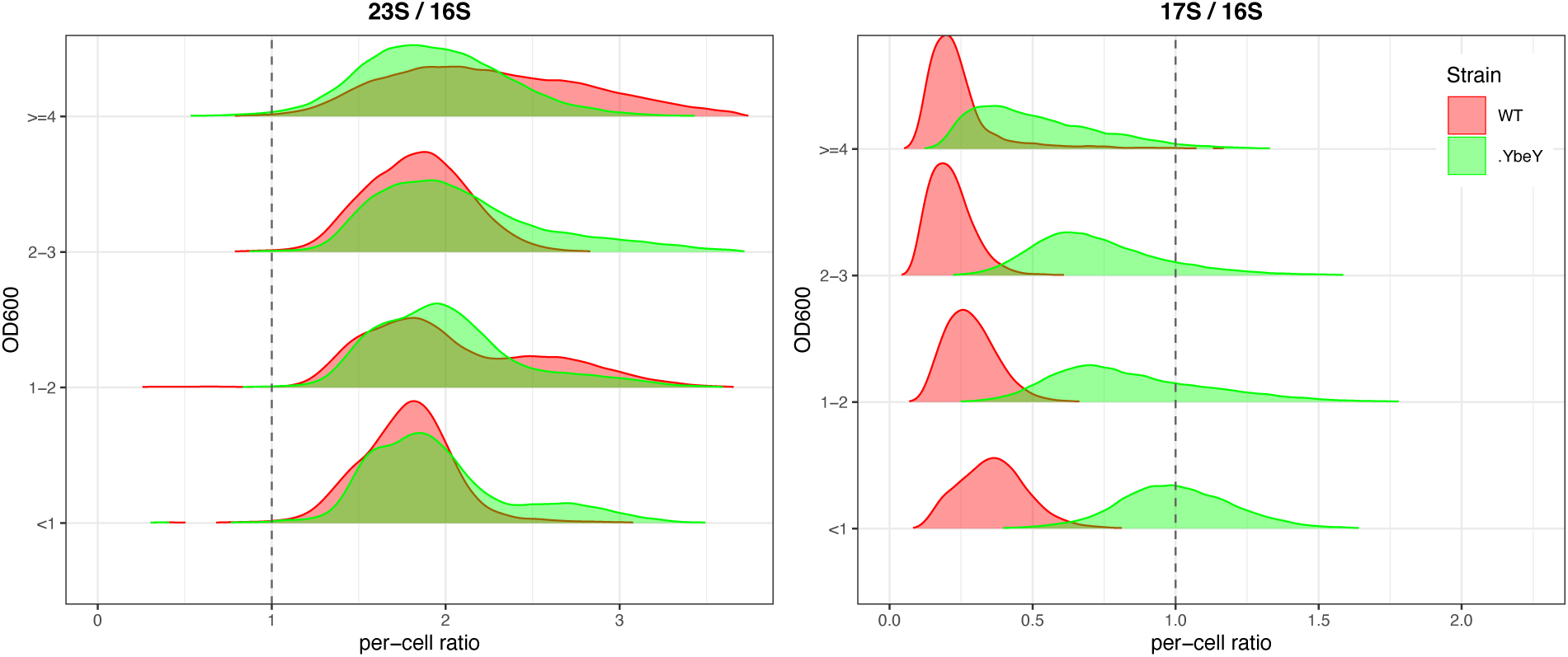
Distribution of per-cell ratios of WT and ΔYbeY 23S/16S and 17S/16S pooled from all experiments.

**Supplementary Figure 13.**
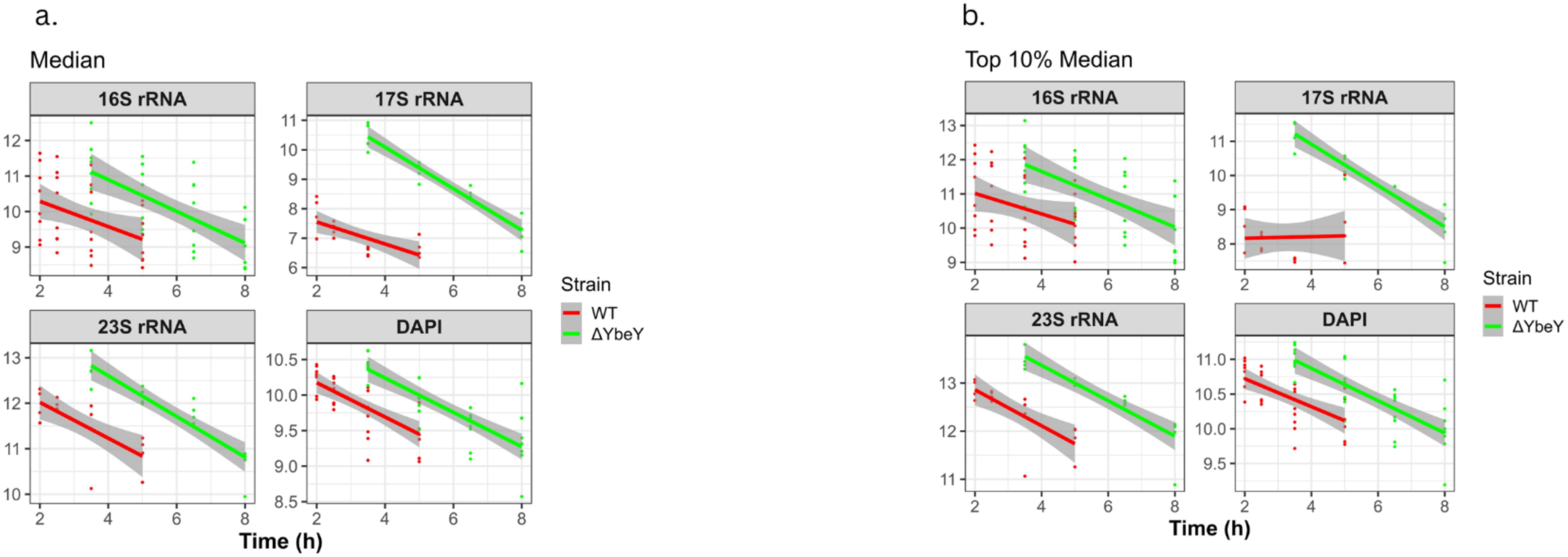
Aggregate-level analysis of the temporal evolution of median rRNA levels and variation. Data points from independent experiments are represented by points (red for WT and green for ΔybeY) and regression lines with shaded 95% CI are colored accordingly.

**Supplementary Figure 14.**
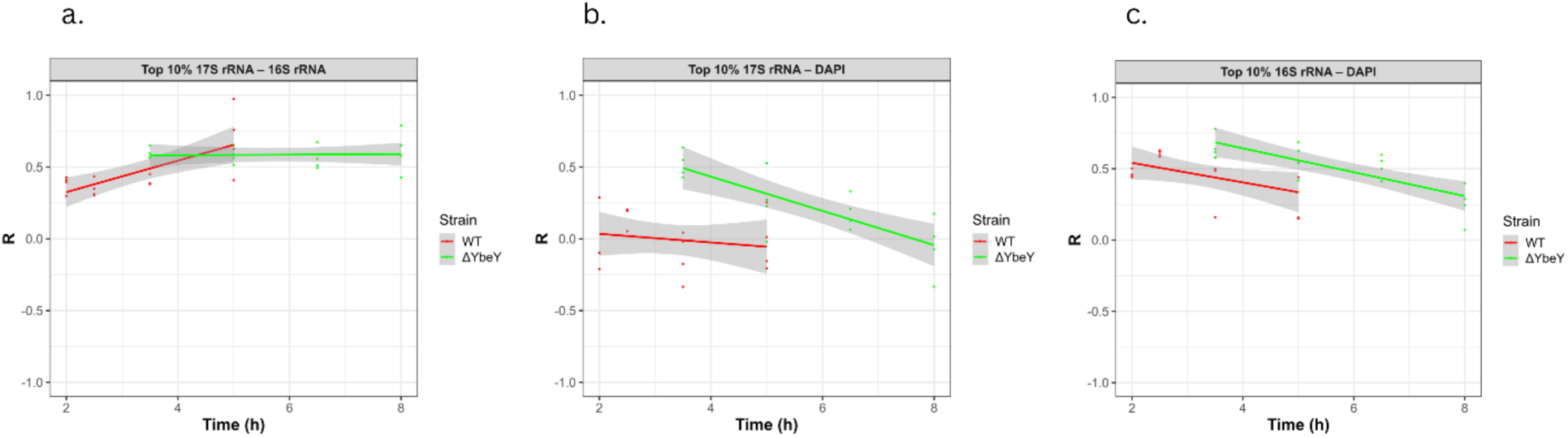
Linear regression model for correlation between WT and ΔybeY cells, manifesting top 10% signal from the second growth culture: a) 17S vs. 16S, b) 17S vs. DAPI, c) 16S vs. DAPI.

**Supplementary Figure 15.**
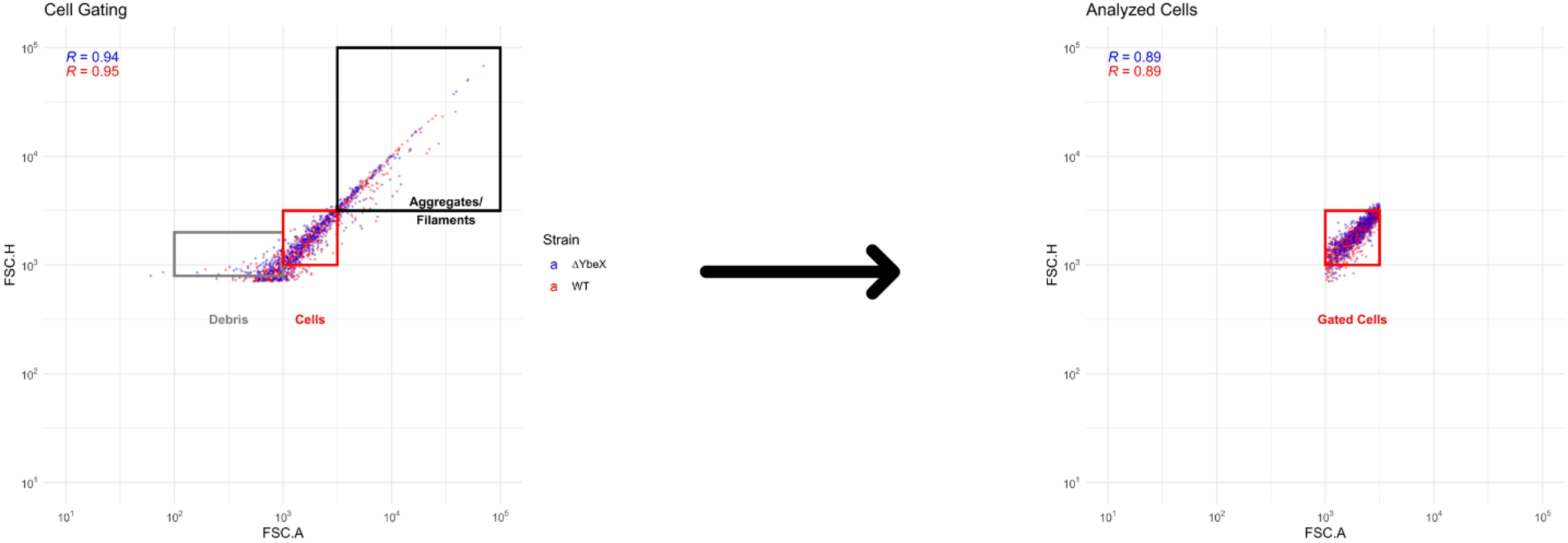
Gating strategy for single-cell flow cytometry analysis. (Left) FSC-A versus FSC-H density plot showing all acquired events for WT (red) and ΔybeX (blue) cultures. Three populations are indicated: debris (low FSC-A, gray box), single cells (intermediate FSC-A/FSC-H with strong linear correlation, red box), and aggregates/filaments (high FSC-A/FSC-H, black box). Pearson correlation coefficients between FSC-A and FSC-H are shown for ΔybeX (R = 0.94, blue) and WT (R = 0.95, red). (Right) The gated singlet population retained for downstream analysis, restricted to the dense core region (10³ ≤ FSC-A, FSC-H ≤ 10³·⁵). Post-gating correlations are R = 0.89 for both strains. Events outside this gate — corresponding to debris, aggregates, filaments, and cells with poor FSC-A/FSC-H correlation — were excluded from all subsequent analyses. Data shown are representative of at least three biological replicates.

## SUPPORTING MATERIALS

### Reagents

**Table 1.**
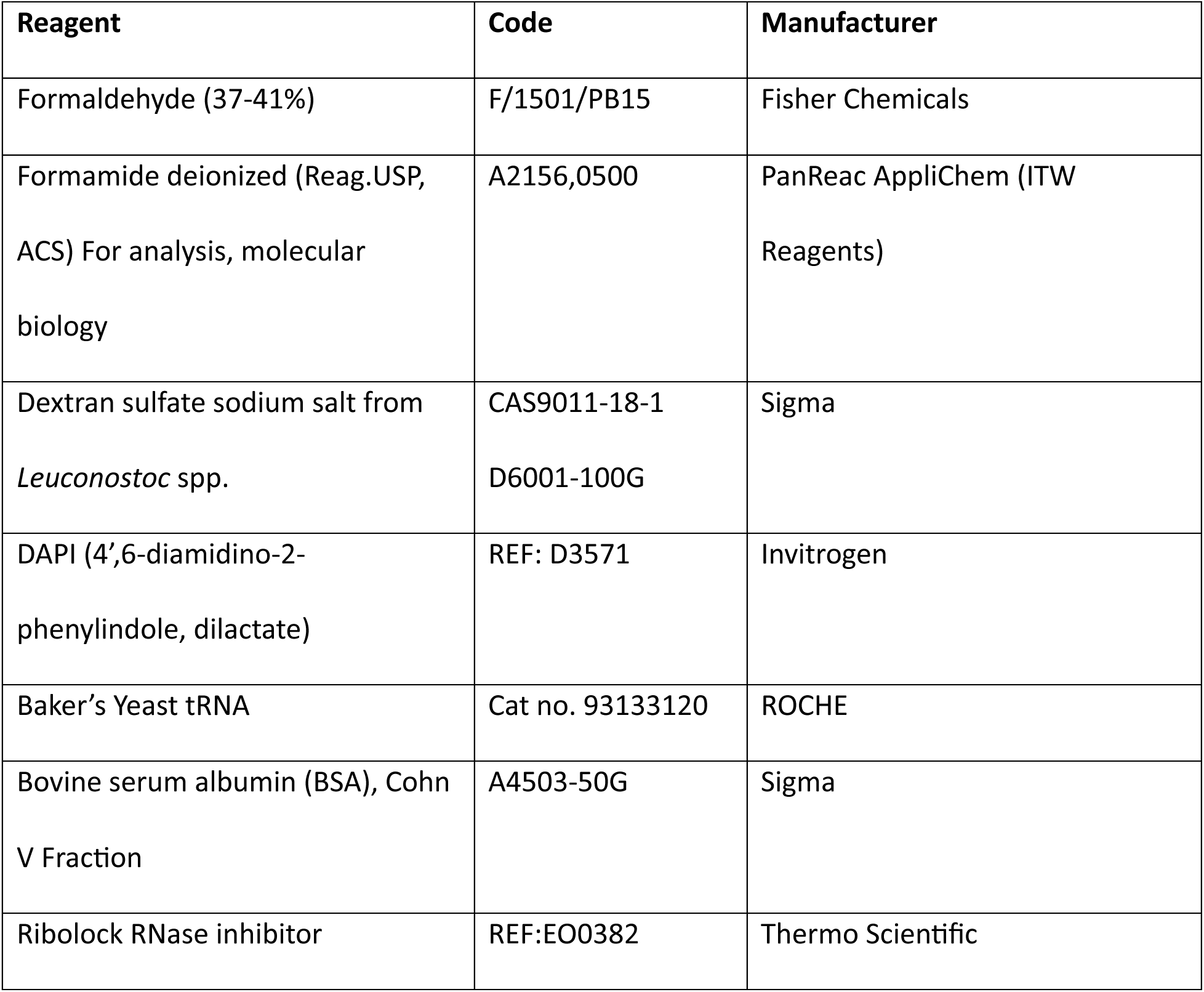
List of reagents used for FISH-FLOW.

**Table 2.**
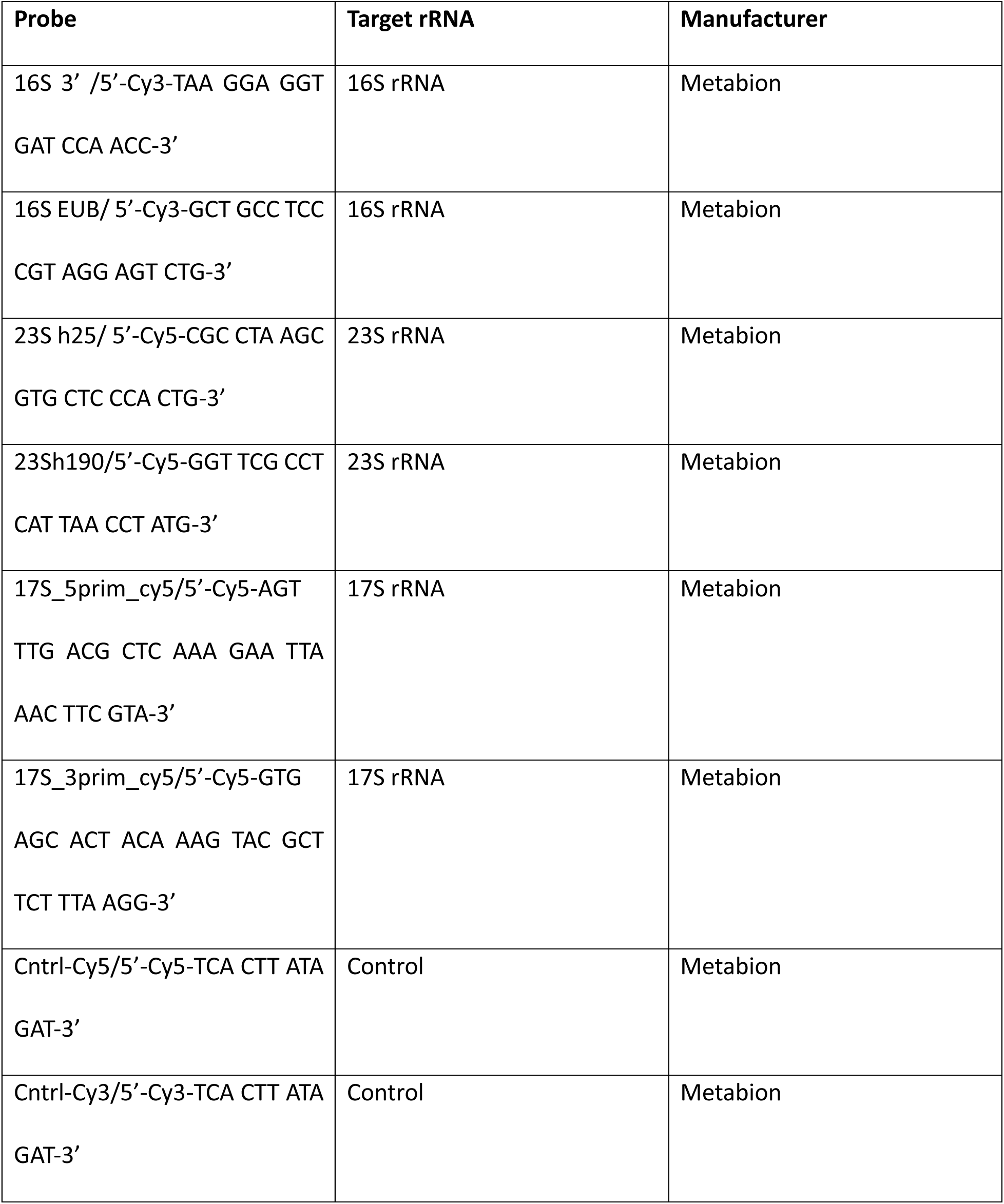
List of probes targeting 23S/16S/17S rRNA.

**Table 3.**
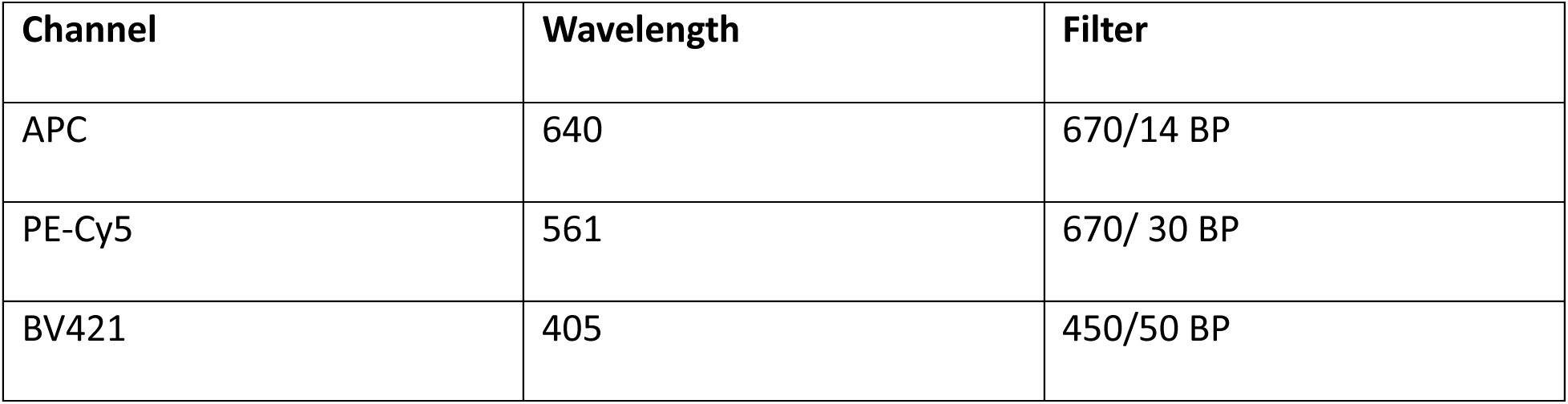
BD LSRFortessa channels for Cy5, Cy3 and DAPI and the respective spectrum range.

