## Supplementary Figures and Tables for "Single-cell analysis of pre-rRNA in *Escherichia coli* indicates distinct pathways of action for YbeX and YbeY proteins in ribosome biogenesis"

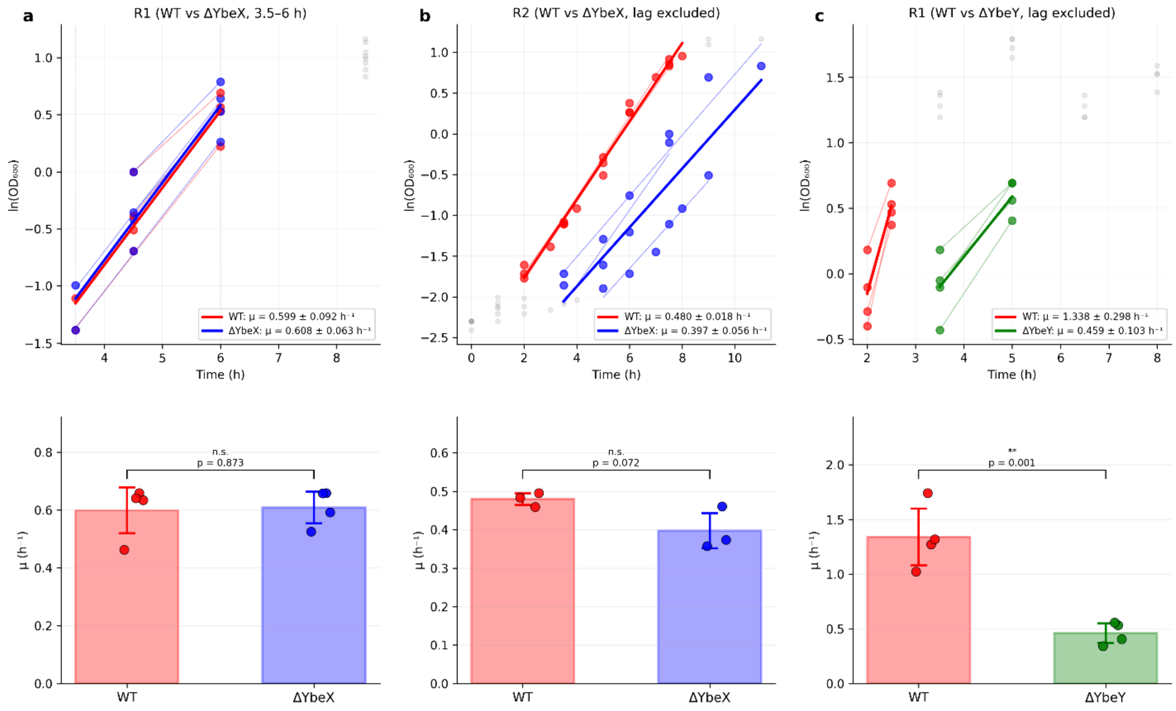


***Supplementary Figure 1.*** *Exponential growth of BW25113, ΔybeX and ΔybeY strains. a) Linear regression of ln($OD_{600}$) against time of wt and ΔybeX in the first growth round. The bar plots show exponential growth rates per hour (μ,$h^{-1}$)). Both strains show similar growth rates (p = 0.87, Student's t-test). b) Linear regression of wt and ΔybeX in the second round of growth. No statistically significant difference in the exponential growth rate was observed between the two strains (p = 0.07). c) Linear regression of wt and ΔybeY, with growth rates shown as bar plots, ΔybeY grew ~3-fold slower than wt (p = 0.001). Data points represent individual biological replicates; error bars indicate SD. Lag-phase and stationary-phase data points were excluded from the regression fits.*

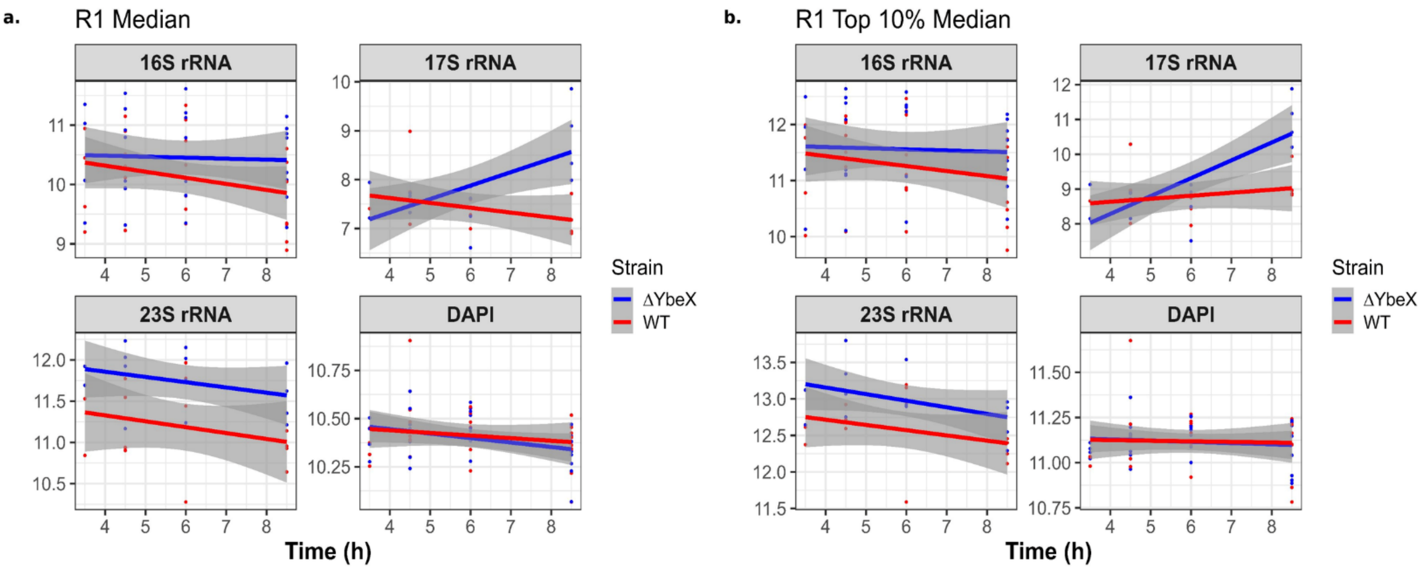


***Supplementary Figure 2.*** *Aggregate-level analysis of all flow cytometry experiments. a. Linear regression of median rRNA and DAPI signals. The median rRNA levels for each independent experiment are plotted against growth time from the start of the experiment (dilution of overnight culture into Mg-limited media). b. Linear regression of interquartile ranges (IQR) of each sample. Data points from independent experiments are represented by points (red for WT and blue for ΔybeX) and regression lines with shaded 95% CI are colored accordingly.*

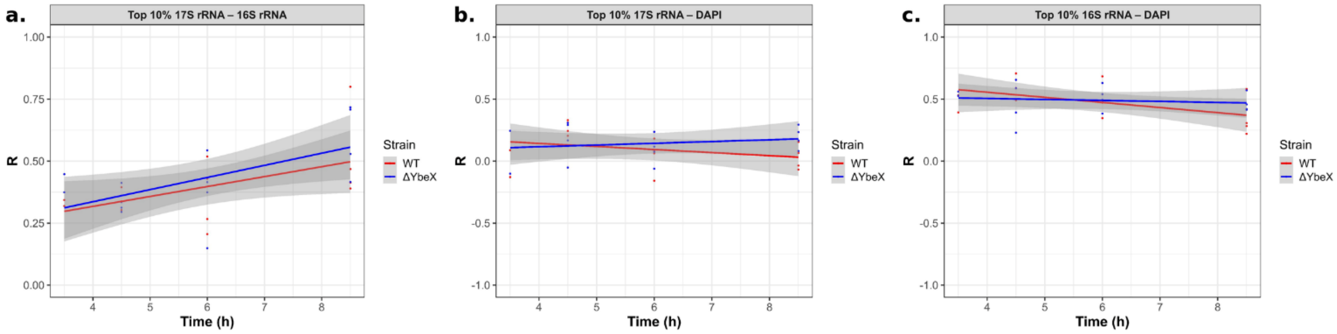


***Supplementary Figure 3.*** *Top 10% (by 17S signal) correlation between a) 17S-16S, b) 17S-DAPI, c) 16S-DAPI. Data points from independent experiments are represented by points (red for WT and blue for ΔybeX) and regression lines with shaded 95% CI are colored accordingly.*

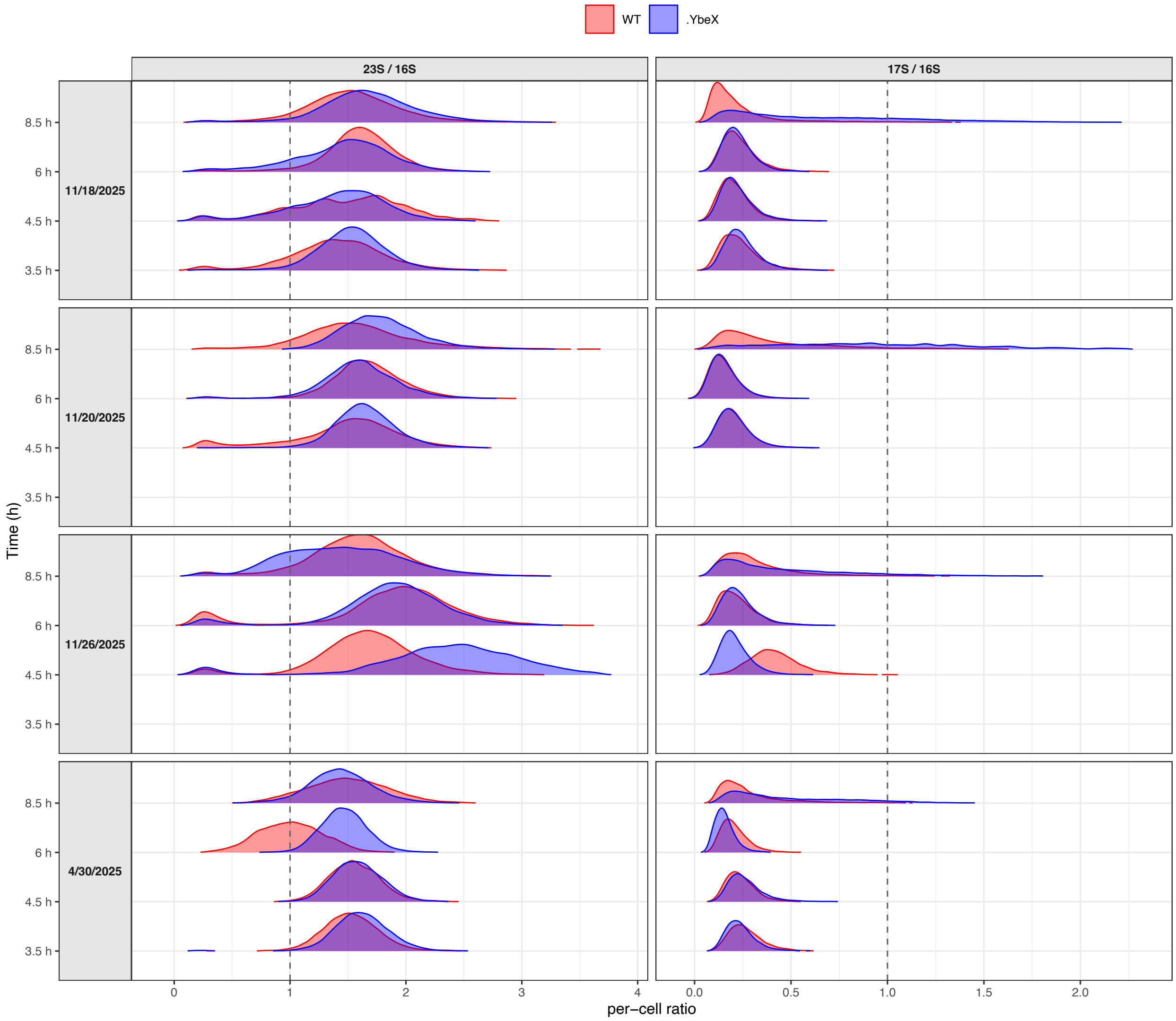


***Supplementary Figure 4.*** *Per-cell 23S/16S and 17S/16S rRNA ratios in WT and ΔybeX, first growth cycle ($Mg^{2+}$-limited MOPS). Linear-scale ridgelines of the per-cell ratio (dashed line = ratio of 1); WT in red, ΔybeX in blue. Each independent experiment in its own panel (faceted by date), time on the y-axis. Note the wide distribution of the ΔybeX 17S/16S at the late time point in every replicate, compared to the narrow 23S/16S. The dates of experiments are shown on the second Y-axis and correspond with those in the supplementary data.*

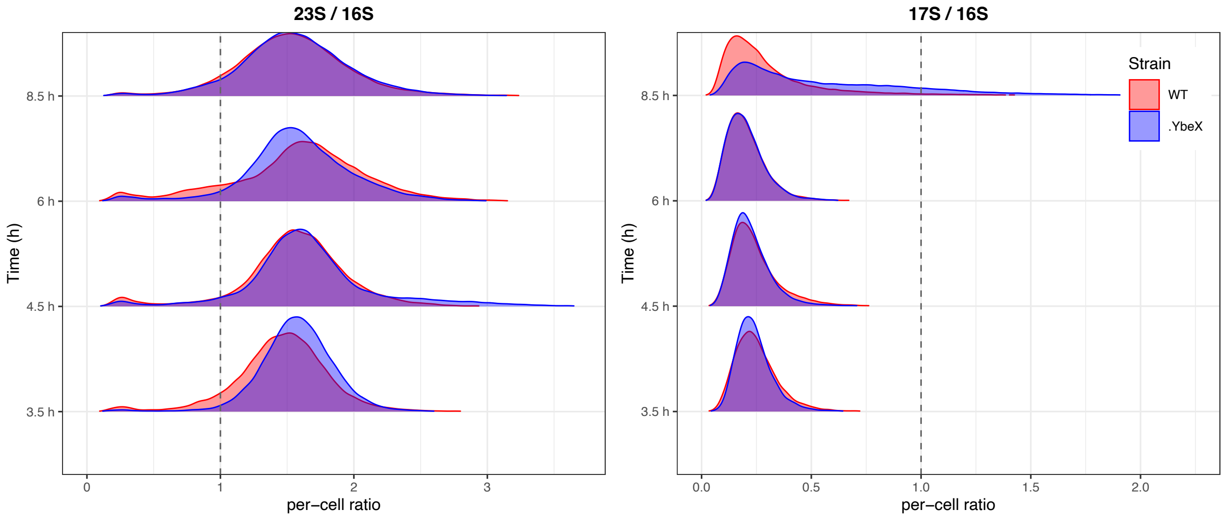


***Supplementary Figure 5.*** *All replicates pooled within each time point. Note the marked broadening of the ΔybeX 17S/16S distribution at the late time point, against the stable, narrow 23S/16S ratio.*

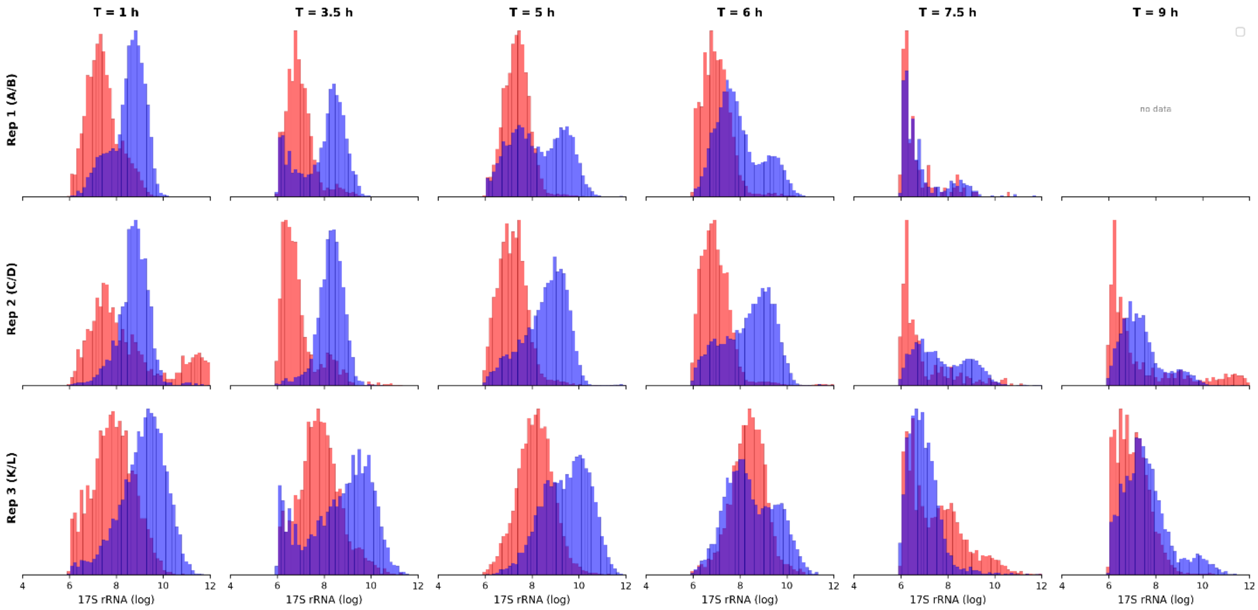


***Supplementary Figure 6.*** *Distribution of 17S rRNA in both BW25113 (red) and ΔybeX (blue) from the second growth round experiment. Three independent experiments are shown. T = 1 h indicates the one hour time point, and so on. The rRNA species is indicated on the X-axis label. All values are in log2 units.*

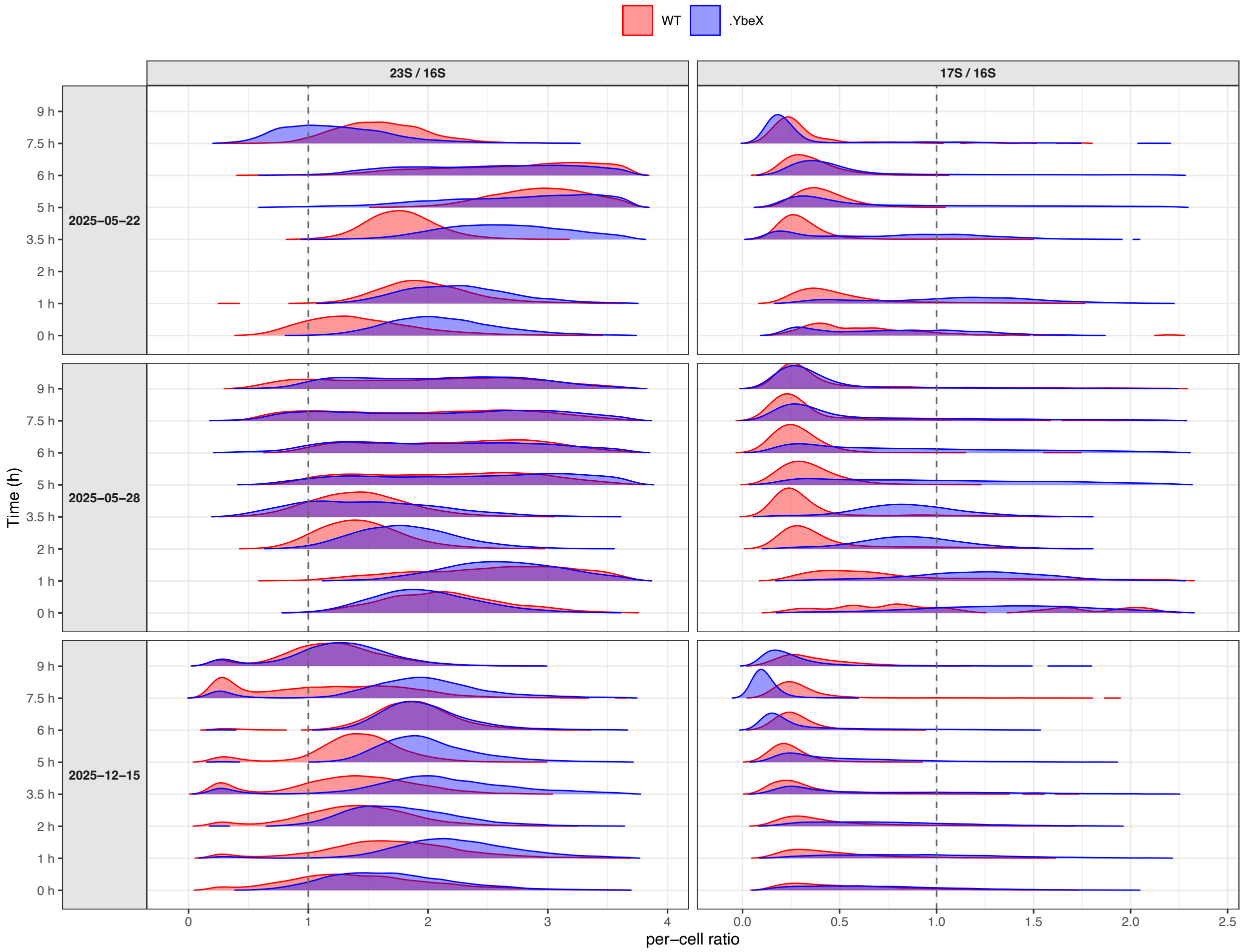


***Supplementary Figure 7.*** *Distributions of per-cell 23S/16S and 17S/16S rRNA ratios in WT and ΔybeX, second growth cycle (outgrowth from stationary phase) The ΔybeX 17S/16S ratio is high and broadly distributed during the lag phase and collapses toward the WT value as visible growth resumes; the 23S/16S ratio stays narrow throughout. The second Y-axis indicates the experiment date, which corresponds to the date in the supplementary data table.*

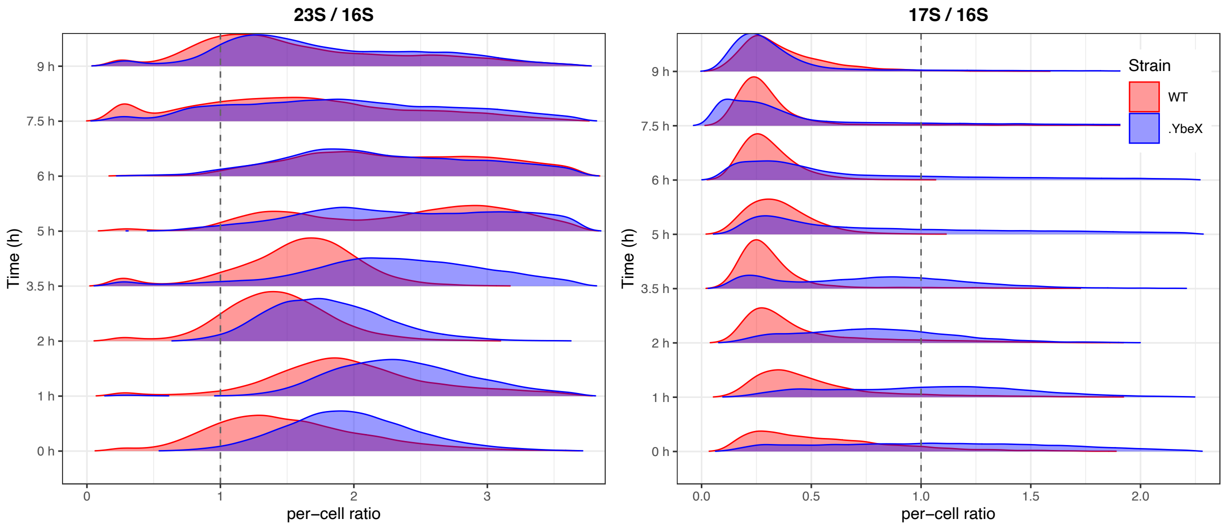


***Supplementary Figure 8.*** *Distribution of per-cell ratios of 23S/16S and 17S/16S rRNAs from the pooled data of the second-round growth experiment.*

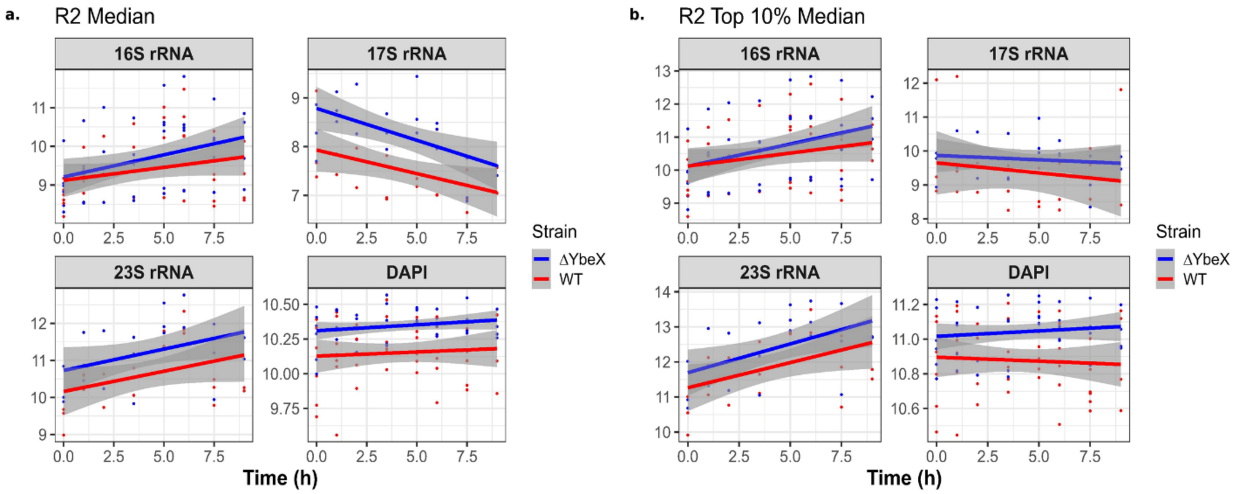


***Supplementary Figure 9.*** *Aggregate-level analysis of the temporal evolution of median rRNA levels and variation in the second growth cycle experiment. Data points from independent experiments are represented by points (red for WT and blue for ΔybeX) and regression lines with shaded 95% CI are colored accordingly. accordingly.*

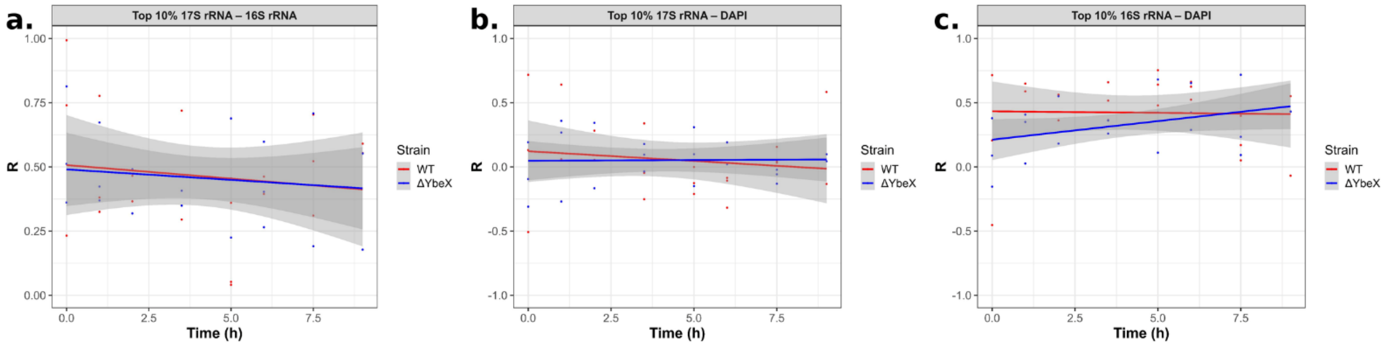


***Supplementary Figure 10.*** *Linear regression model for correlation between WT and ΔybeX cells manifesting top 10% signal from the second growth culture: a) 17S vs 16S, b) 17S vs DAPI, c) 16S vs DAPI.*

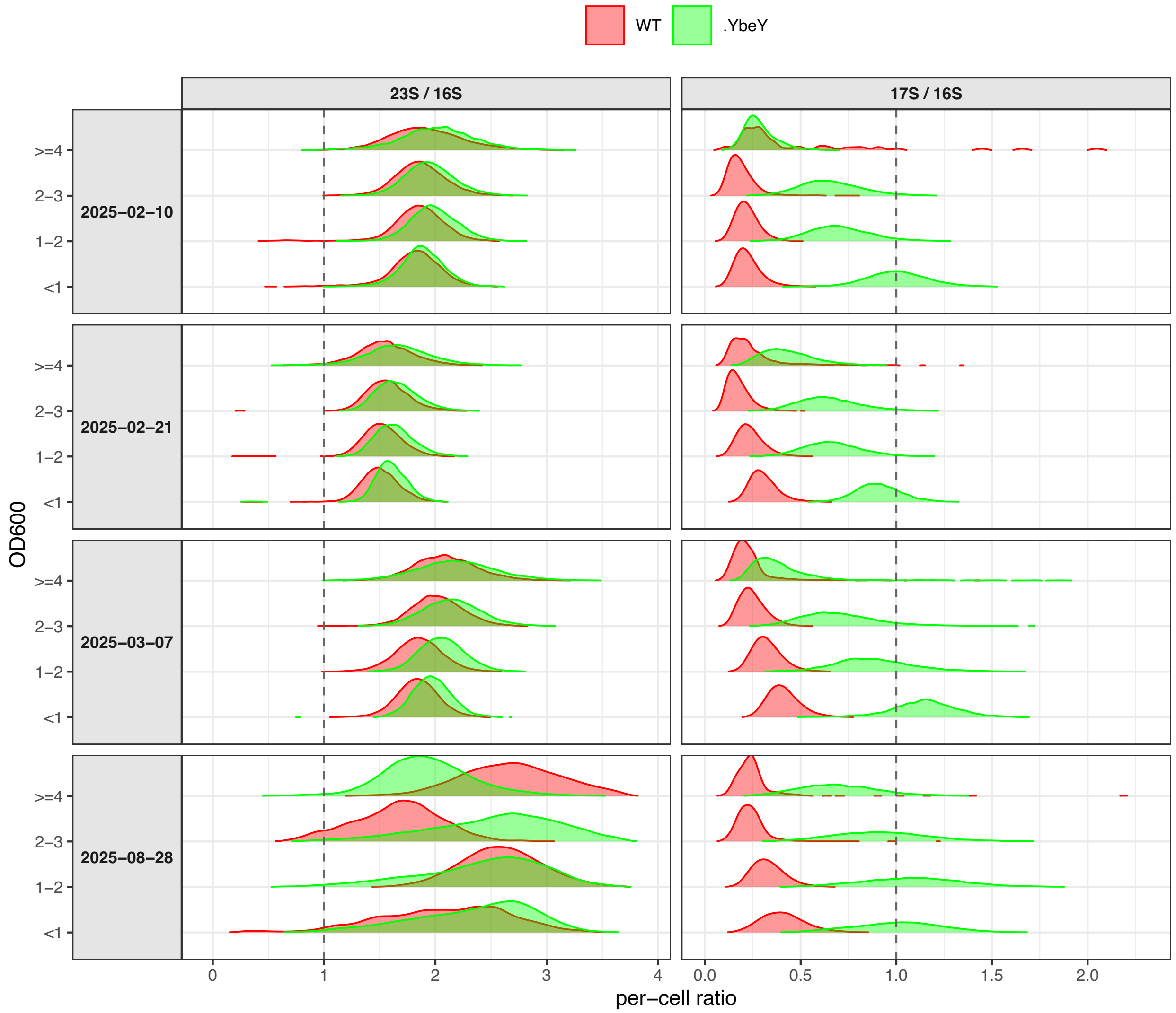


***Supplementary Figure 11.*** *Distribution of per-cell 23S/16S and 17S/16S rRNA ratios in WT and ΔybeY strains, at indicated* $OD_{600}$ *values. Layout is as in Supplementary Figure 4 with ΔybeY shown in green and* $OD_{600}$ *on the y-axis. The ΔybeY 17S/16S ratio is uniformly elevated as a narrow, unimodal distribution, in contrast to the heterogeneous broadening seen in ΔybeX.*

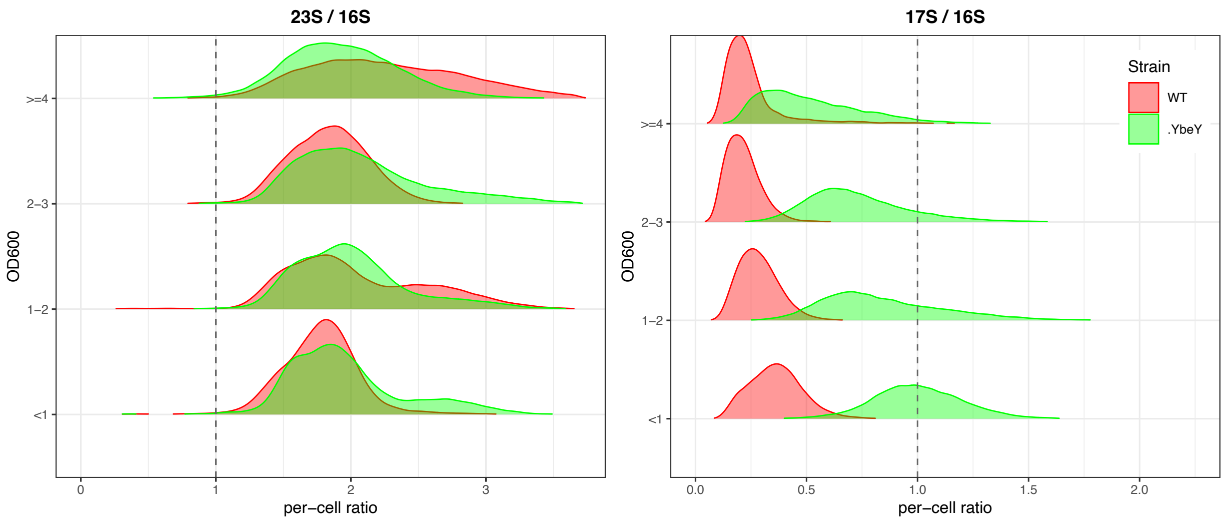


***Supplementary Figure 12.*** *Distribution of per-cell ratios of WT and ΔYbeY 23S/16S and 17S/16S pooled from all experiments.*

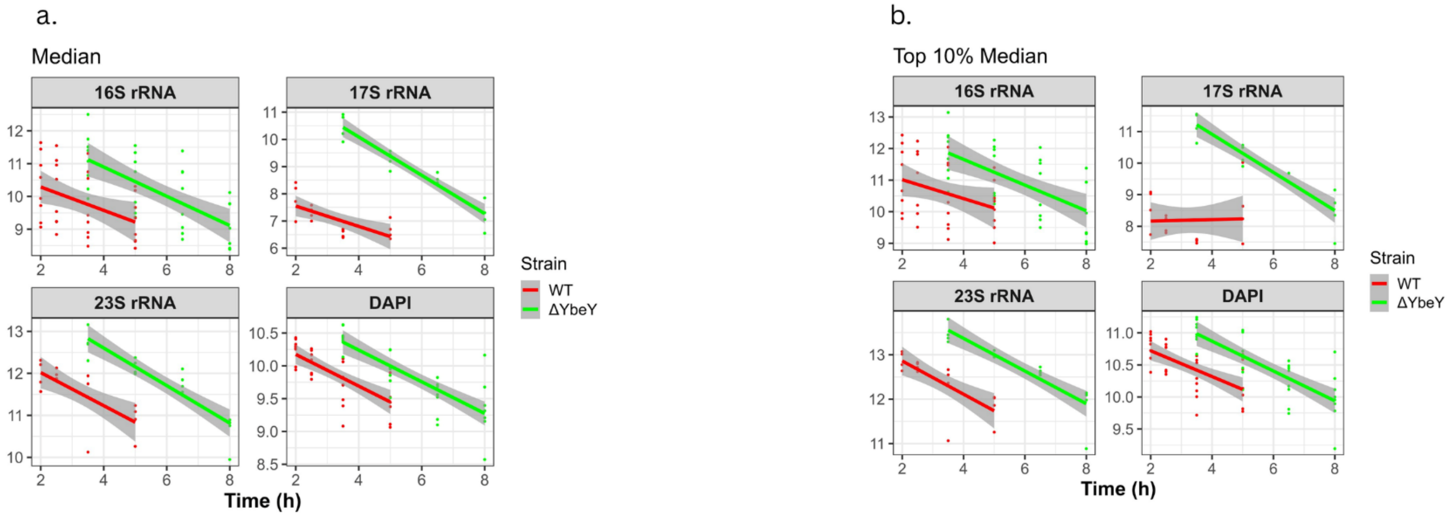


***Supplementary Figure 13.*** *Aggregate-level analysis of the temporal evolution of median rRNA levels and variation. Data points from independent experiments are represented by points (red for WT and green for ΔybeY) and regression lines with shaded 95% CI are colored accordingly.*

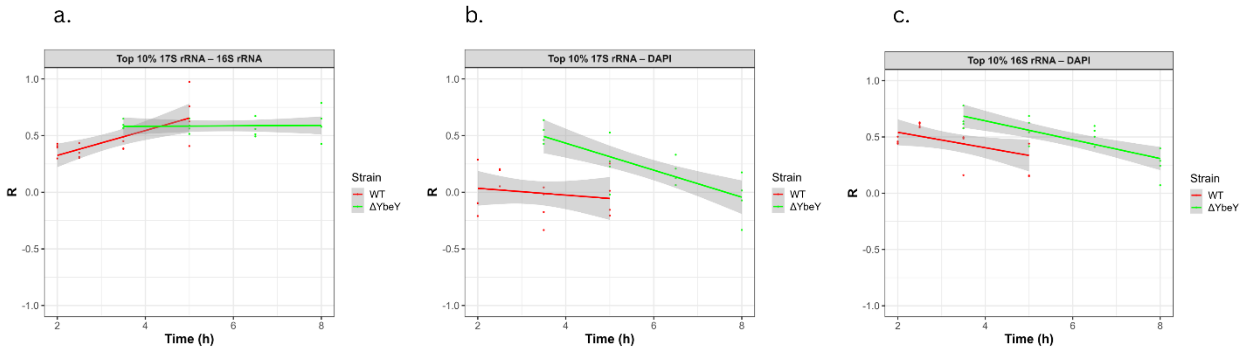


***Supplementary Figure 14.*** *Linear regression model for correlation between WT and ΔybeY cells, manifesting top 10% signal from the second growth culture: a) 17S vs. 16S, b) 17S vs. DAPI, c) 16S vs. DAPI.*

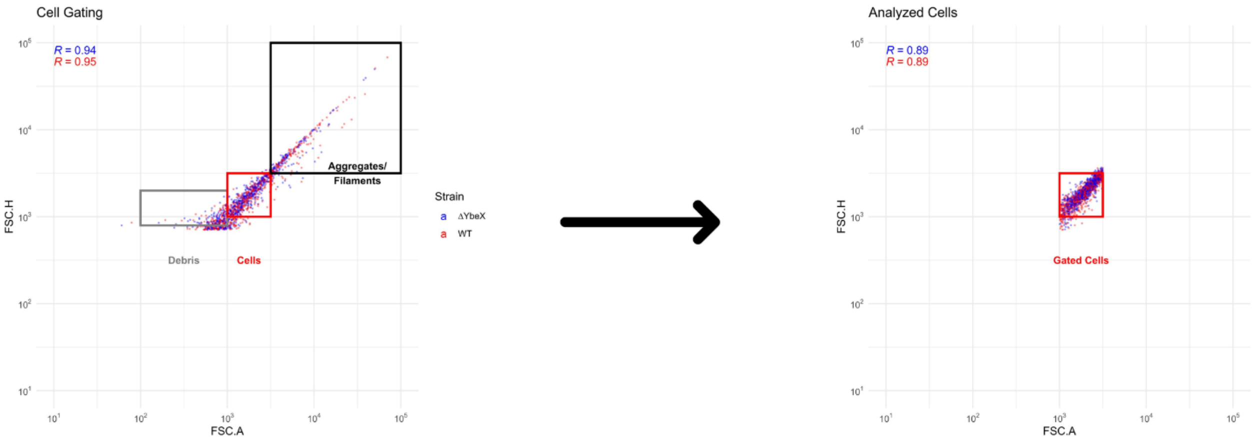


***Supplementary Figure 15.*** *Gating strategy for single-cell flow cytometry analysis. (Left) FSC-A versus FSC-H density plot showing all acquired events for WT (red) and ΔybeX (blue) cultures. Three populations are indicated: debris (low FSC-A, gray box), single cells (intermediate FSC-A/FSC-H with strong linear correlation, red box), and aggregates/filaments (high FSC-A/FSC-H, black box). Pearson correlation coefficients between FSC-A and FSC-H are shown for ΔybeX (R = 0.94, blue) and WT (R = 0.95, red). (Right) The gated singlet population retained for downstream analysis, restricted to the dense core region ($10^3$ ≤ FSC-A, FSC-H ≤ $10^{3.5}$). Post-gating correlations are R = 0.89 for both strains. Events outside this gate — corresponding to debris, aggregates, filaments, and cells with poor FSC-A/FSC-H correlation — were excluded from all subsequent analyses. Data shown are representative of at least three biological replicates.*

**SUPPORTING MATERIALS:**

**Reagents**

**Table 1.** List of reagents used for FISH-FLOW.

| **Reagent** | **Code** | **Manufacturer** |
|---|---|---|
| Formaldehyde (37-41%) | F/1501/PB15 | Fisher Chemicals |
| Formamide deionized (Reag.USP, ACS) For analysis, molecular biology | A2156,0500 | PanReac AppliChem (ITW Reagents) |
| Dextran sulfate sodium salt from *Leuconostoc* spp. | CAS9011-18-1 D6001-100G | Sigma |
| DAPI (4',6-diamidino-2-phenylindole, dilactate) | REF: D3571 | Invitrogen |
| Baker's Yeast tRNA | Cat no. 93133120 | ROCHE |
| Bovine serum albumin (BSA), Cohn V Fraction | A4503-50G | Sigma |
| Ribolock RNase inhibitor | REF:EO0382 | Thermo Scientific |

**Table 2.** List of probes targeting 23S/16S/17S rRNA.

| Probe | Target rRNA | Manufacturer |
|---|---|---|
| 16S 3' /5'-Cy3-TAA GGA GGT GAT CCA ACC-3' | 16S rRNA | Metabion |
| 16S EUB/ 5'-Cy3-GCT GCC TCC CGT AGG AGT CTG-3' | 16S rRNA | Metabion |
| 23S h25/ 5'-Cy5-CGC CTA AGC GTG CTC CCA CTG-3' | 23S rRNA | Metabion |
| 23Sh190/5'-Cy5-GGT TCG CCT CAT TAA CCT ATG-3' | 23S rRNA | Metabion |
| 17S_5prim_cy5/5'-Cy5-AGT TTG ACG CTC AAA GAA TTA AAC TTC GTA-3' | 17S rRNA | Metabion |
| 17S_3prim_cy5/5'-Cy5-GTG AGC ACT ACA AAG TAC GCT TCT TTA AGG-3' | 17S rRNA | Metabion |
| Cntrl-Cy5/5'-Cy5-TCA CTT ATA GAT-3' | Control | Metabion |
| Cntrl-Cy3/5'-Cy3-TCA CTT ATA GAT-3' | Control | Metabion |

**Table 3.** BD LSRFortessa channels for Cy5, Cy3 and DAPI and the respective spectrum range.

| Channel | Wavelength | Filter |
|---|---|---|
| APC | 640 | 670/14 BP |
| PE-Cy5 | 561 | 670/ 30 BP |
| BV421 | 405 | 450/50 BP |
